# Convergent selection in antibody repertoires is revealed by deep learning

**DOI:** 10.1101/2020.02.25.965673

**Authors:** Simon Friedensohn, Daniel Neumeier, Tarik A Khan, Lucia Csepregi, Cristina Parola, Arthur R Gorter de Vries, Lena Erlach, Derek M Mason, Sai T Reddy

**Affiliations:** Department of Biosystems Science and Engineering, ETH Zürich, 4058 Basel, Switzerland

## Abstract

Adaptive immunity is driven by the ability of lymphocytes to undergo V(D)J recombination and generate a highly diverse set of immune receptors (B cell receptors/secreted antibodies and T cell receptors) and their subsequent clonal selection and expansion upon molecular recognition of foreign antigens. These principles lead to remarkable, unique and dynamic immune receptor repertoires^1^. Deep sequencing provides increasing evidence for the presence of commonly shared (convergent) receptors across individual organisms within one species^2-4^. Convergent selection of specific receptors towards various antigens offers one explanation for these findings. For example, single cases of convergence have been reported in antibody repertoires of viral infection or allergy^5-8^. Recent studies demonstrate that convergent selection of sequence motifs within T cell receptor (TCR) repertoires can be identified on an even wider scale^9,10^. Here we report that there is extensive convergent selection in antibody repertoires of mice for a range of protein antigens and immunization conditions. We employed a deep learning approach utilizing variational autoencoders (VAEs) to model the underlying process of B cell receptor (BCR) recombination and assume that the data generation follows a Gaussian mixture model (GMM) in latent space. This provides both a latent embedding and cluster labels that group similar sequences, thus enabling the discovery of a multitude of convergent, antigen-associated sequence patterns. Using a linear, one-versus-all support vector machine (SVM), we confirm that the identified sequence patterns are predictive of antigenic exposure and outperform predictions based on the occurrence of public clones. Recombinant expression of both natural and *in silico*-generated antibodies possessing convergent patterns confirms their binding specificity to target antigens. Our work highlights to which extent convergence in antibody repertoires can occur and shows how deep learning can be applied for immunodiagnostics and antibody discovery and engineering.

## RESULTS

Targeted deep sequencing of the rearranged BCR locus reveals the antibody repertoire of B cells in a given tissue or cell population^11^. Here we used deep sequencing to analyse the antibody repertoires in the bone marrow of 45 BALB/c mice, which were divided into four separate cohorts immunized with protein antigens of either ovalbumin (OVA), hen egg lysozyme (HEL), blue carrier protein (BCP) or respiratory syncytial virus fusion protein (RSV-F). OVA, HEL and BCP cohorts were further subdivided into groups receiving zero, one, two or three booster immunizations; serum enzyme-linked immunoabsorbance assays (ELISA) confirmed antigen-specific antibody titers in all mice, with mice receiving only a primary immunization exhibiting substantially weaker titers (**Supplementary Table 1**). RNA was extracted in bulk from bone marrow cells and variable heavy (V_H_) chain IgG sequencing libraries were prepared using a two-step RT-PCR protocol, which included molecular barcoding for error and bias correction^12^. Libraries were sequenced on an Illumina MiSeq, quality processed and aligned, yielding across all mice a total of 243’374 unique combinations of all three V_H_ complementarity-determining regions (CDRHs) (**Supplementary Fig. 1a, b**). Similar to previous studies^3^, we observed the occurrence of public clones (defined here as identical CDRH1-CDRH2-CDRH3 amino acid (a.a.) sequences that occur in more than one mouse). However, the occurrence of public clones was not limited by antigen exposure, as on average 16% of the sequences found in a given repertoire were shared with repertoires outside the respective antigen cohort, whereas only ∼4% were limited to a specific antigen cohort (**Supplementary Fig. 1c**).

To evaluate to which extent convergent selection occurs that is beyond the occurrence of public clones, we developed a deep learning workflow which utilizes CDRH1, CDRH2 and CDRH3 and their appropriate sequence combinations as input to a VAE model. Deep generative models using variational inference have recently been used for TCR repertoire modelling and protein fitness prediction^13-15^. Unlike this previous work however, we assume that sequences in latent space are generated by a GMM, which enables the VAE to estimate the densities of different clonal lineages and signatures more accurately (**Fig. 1**). The VAE consists of deep neural networks which are used to encode and decode sequences and are optimized with respect to the GMM prior and their ability to reconstruct the input sequences (**Extended Data Fig. 1)**, with similar sequences falling into the same cluster and closely related clusters occupying similar regions in latent space (**see Methods**). Increasing the dimensionality of the latent encoding and the number of clusters improved the reconstruction ability of the model; by using a ten-dimensional latent space with 2,000 clusters, we were able to achieve a reconstruction accuracy of 93.4% (**Extended Data Fig. 2**). We used principal component analysis (PCA) to visualize the latent encodings and found that highly similar sequences did indeed map to the same cluster and areas within the latent space (**Fig. 2a**). The VAE model also captured important information such as CDRH3 length and variable germline segment (V-gene), whereas similar V-gene families were mapped into similar areas of the latent space (**Extended Data Fig. 3**). While visual inspection revealed areas of possible antigen-associated sequence convergence (**Fig. 2a, Extended Data Fig. 4**), we aimed to quantify the amount of convergence by identifying latent clusters that were significantly associated for each respective antigen, and whether these convergent sequences could be used to predict the antigen exposure of a given mouse. Sequences were grouped into their respective clusters and the recoded repertoires were used to train and cross-validate a linear, one-versus-all SVM classifier of antigen exposure (**Supplementary Fig. 2**). It is important to note that the VAE allows cluster assignments of unseen data and thus an independent embedding could be trained for each fold. In order to establish a baseline for this workflow, we also trained a linear SVM on the occurrence of public clones (exact CDRH1-CDRH2-CDRH3 a.a. sequence matches), which yielded an accuracy of 42% for prediction of antigen exposure (5-fold cross-validation). In contrast, when using VAE-based cluster assignments and subsequently encoding repertoires based on cluster enrichment, the resulting classifiers were able to achieve a prediction accuracy of over 80% (**Fig. 2b, Extended Data Fig. 5 and 6**). We then performed a simple, non-parametric permutation test for each cluster and each cohort at a significance level of α=0.05. Bonferroni correction was conducted in order to account for multiple testing, yielding 60 (BCP, 6’664 sequences), 61 (RSV-F, 7’064 sequences), 68 (OVA, 7’389 sequences) and 73 (HEL, 9’628 sequences) significantly enriched convergent antibody clusters in each cohort. While the exact number of convergent clusters and sequences slightly varies with the number of latent space clusters and the initialization of the VAE, the anti-correlation between protein antigen size and complexity and identified sequences suggests that convergence may be driven by the presence of a few immunodominant epitopes. Closer inspection revealed that not every mouse expressed all of their respective convergent clusters, but rather a smaller, yet still predictive subset (**Fig. 2c**). Furthermore, mice that only received a primary immunization without any subsequent booster immunization also exhibited a decreased enrichment of convergent clones (**Fig. 2c**, area between dashed red lines), a finding that correlates well with the measured serum titters (**Supplementary Table 1).** Example logos generated by sequences mapping into the same cluster visualize how the VAE model is able to identify diverse and biologically meaningful groupings (**Fig. 2d**). Furthermore, comparing aggregated sequence logos to those generated from single repertoires shows the potential diversity of the convergent sequence space and highlights that convergence is not limited only to highly similar, public CDRHs (**Extended Data Fig. 7**). As an additional, yet simple example of convergence, we observed a frequently occurring asparagine (N) residue in the first position of CDRH3 of multiple RSV-F-associated clusters (**Fig. 2d**).

**Figure 1.**
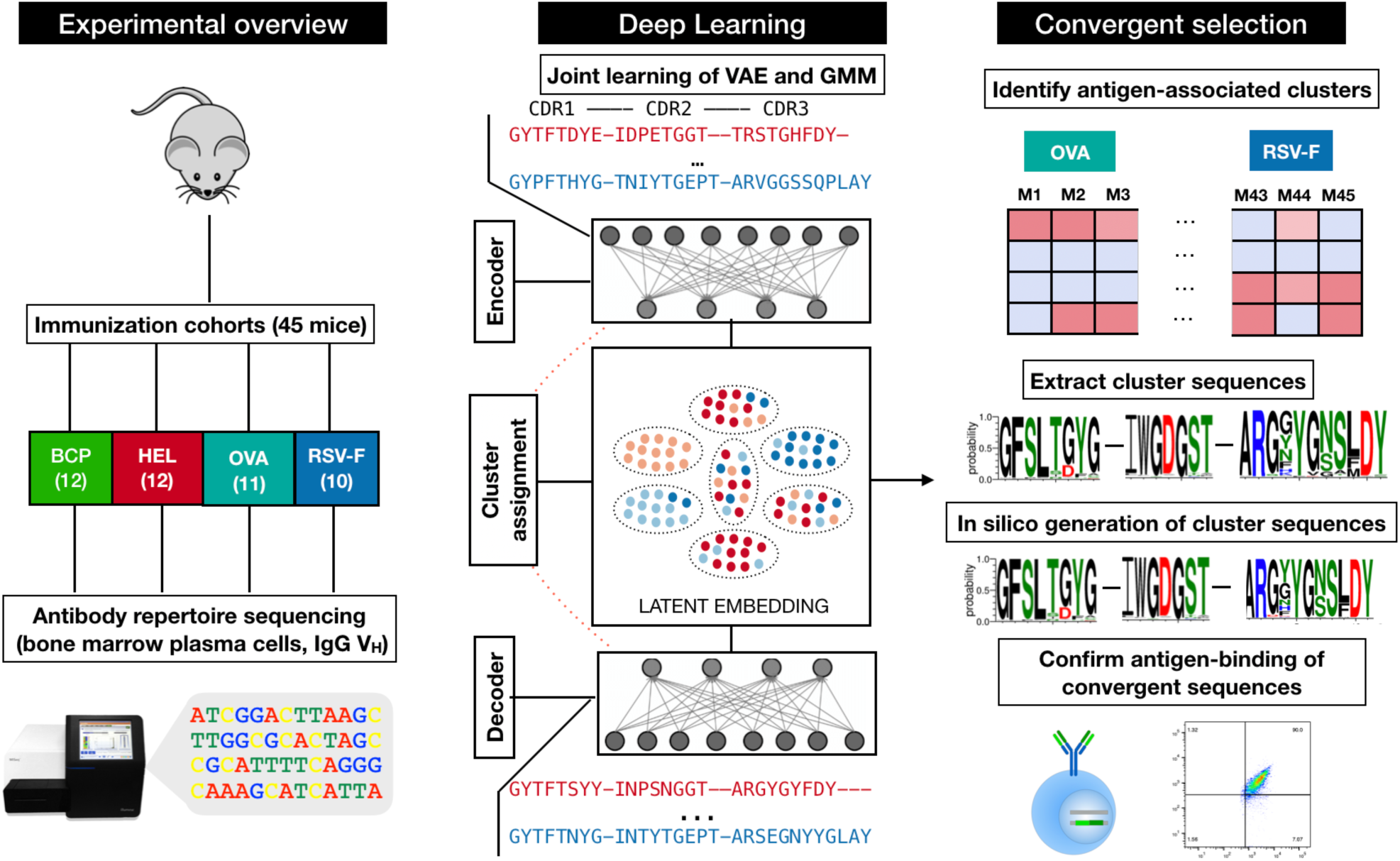
Schematic overview of deep learning on antibody repertoires of immunized mice. Antibody repertoires from the bone marrow of mice immunized with various antigens are deep sequenced. Antibody sequences are then used to train a variational autoencoder (VAE) with a Gaussian mixture model (GMM) clustering of the latent space. The VAE model is able to both generate novel sequences and assign input sequences to distinct clusters based on their latent space embedding. Cluster assignments are used to identify convergent sequences that are heavily enriched in a specific repertoire or antigen cohort. Natural and in-silico generated sequences from antigen-associated clusters are expressed as full-length IgG and verified as binding antibodies.

**Figure 2.**
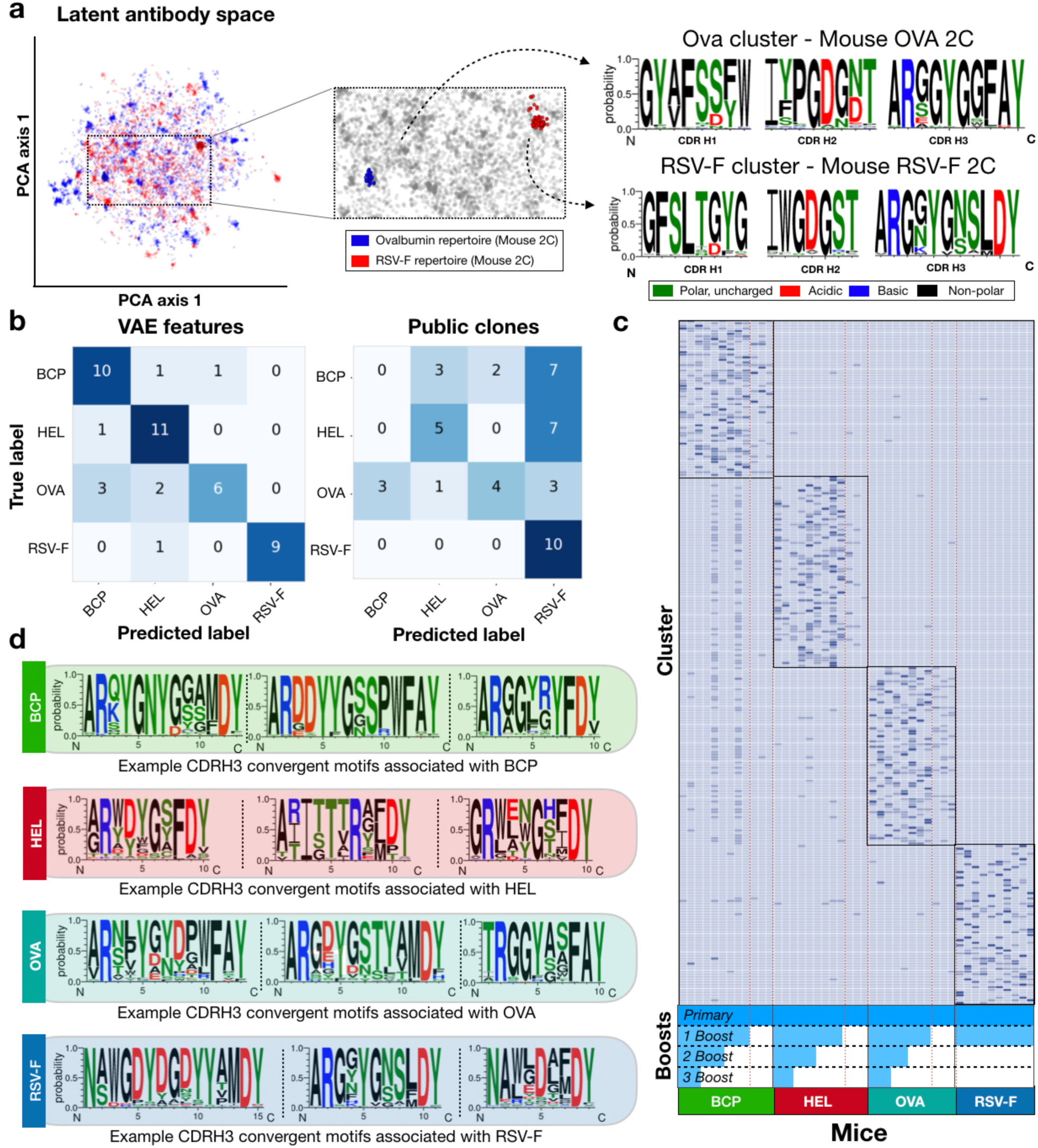
Identification and characterization of convergent antigen-associated sequences. a, Ten-dimensional latent space of two antibody repertoires visualized by principal component analysis (PCA). Blue and red dots indicate sequences belonging to one OVA (2C) and RSV-F (2C) repertoire, respectively. Enlarged area highlights two learned clusters only containing sequences specific to one repertoire and their respective sequence motifs. b, Antibody repertoires are transformed into vectors based on the learned sequence clusters in latent space. Recoded vectors are used as input for a linear support vector machine (SVM) classifier of antigen exposure. Confusion matrices show the aggregated prediction results of each model during 5-fold cross-validation using the cluster labels and raw sequences as features. c, Heatmap contains all predictive and convergent sequence clusters for each cohort. Dashed red line indicates mice that only received the primary immunization. d, Example sequence logos of convergent clusters found in each antigen cohort.

In order to confirm that antigen-associated sequence convergence corresponds to antigen binding, we utilised a mammalian cell (hybridoma) antibody surface display and secretion system coupled to CRISPR-Cas9-mediated library integration^16^. An antibody library was generated using a small subset of convergent V_H_ sequences derived from different clusters combined with a variable light chain (V_L_) library cloned from cohort-matched mouse repertoires (**Extended Data Fig. 8**). ELISAs performed on the supernatant of the library cell lines demonstrated that these convergent pools possessed cognate antigen specificity (**Fig. 3a**). We then proceeded to more closely investigate V_H_ variants from the OVA and RSV-F pools through single clone isolation by fluorescence-activated cell sorting (FACS) (**Supplementary. Fig. 3**). The antigen-specific binding of monoclonal cell lines was confirmed by flow cytometry (**Fig. 3b**) and ELISA (**Supplementary Fig. 4**) and their V_H_ chains were identified by sequencing (**Supplementary Fig. 5**). This procedure allowed us to confirm antigen specificity of 6 (out of 6 selected) OVA and 3 (out of 4 selected) RSV-F convergent V_H_ sequences (**Extended Data Table 1**). V_H_ chains were able to pair with V_L_ chains from a different mouse repertoire, additionally highlighting convergence with respect to V_H_ chain-dominated binding (**Supplementary Tables 2-5**). While all antigens were associated with a variety of V-gene germlines, we noticed that convergent antibodies were utilizing different V-gene segments in an antigen-dependent manner, highlighting that the original V-gene germline contributes to convergent selection (**Fig. 3f, Extended Data Fig. 9).**

**Figure 3.**
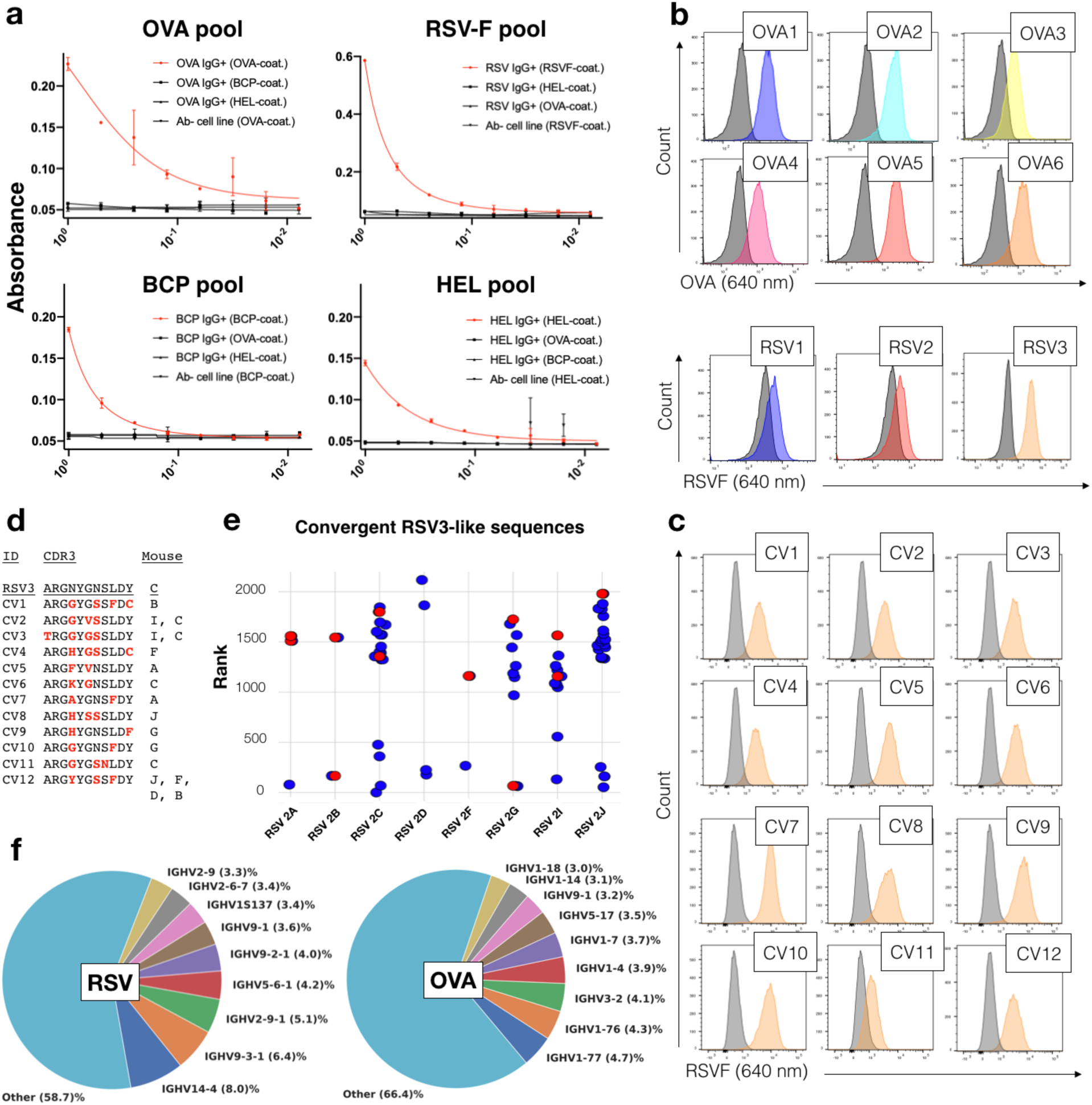
Convergent clusters contain antigen-specific antibodies. a, Dose-dependent absorbance curves of supernatant prepared from hybridoma cells expressing antibodies with convergent variable heavy (V_H_) chain pools for each antigen. b, Flow cytometry histograms of six monoclonal cell populations each utilizing a different convergent OVA-associated or RSV-F associated V_H_. Grey histograms represent negative controls, colored histograms show the convergent antibodies. c, Flow cytometry histograms of 12 monoclonal cell populations of convergent variants (CV), which use a different V_H_ sequence from the same cluster as RSV3. d, Table shows the CDRH3s of the selected CVs and the RSV-F immunized mouse repertoire in which they were found. Red letters indicate differences to the initially discovered sequence RSV3 sequence. e, Scatterplot shows the frequency-rank distributions per mouse repertoire of CVs from RSV3 cluster. Red dots highlight V_H_ confirmed to be binding in **c**. e, Pie charts show the nine most utilized V-gene germlines in convergent clones for both RSV-F and OVA.

In order to confirm that antibody sequence variants mapping to the same convergent cluster were also antigen-specific, we recombinantly expressed 12 convergent V_H_ variants (derived from other mice immunized with the same antigen) from the cluster mapping to one of the confirmed RSV-F binders (RSV3, **Supplementary Fig. 6**). These 12 convergent variants were expressed with the same V_L_ of RSV3. Flow cytometry confirmed that all 12 of these convergent variants were indeed antigen-specific (**Fig. 3c**). Using standard clonotype definitions of 100% or 80% V_H_ CDRH3 a.a. identity ^2,4^, only zero or five of the 12 variants, respectively, would have been identified as convergent across repertoires (**Fig. 3d**). In contrast, the VAE model was able to discover variants of RSV3 with as low as 64% CDRH3 a.a. identity (4 out of 11 mismatches), verifying the large potential diversity revealed by the previous logo plots (**Fig. 2d, Fig. 3f**). Besides their sequence diversity, these clones also confirmed the large abundance range with confirmed binders being of high, medium and low frequencies in their respective mouse repertoires (**Fig. 3e**).

Finally, we aimed to understand how well the VAE model is able to generalise to unseen data. To start, we experimentally produced an antibody CDRH3 library of the RSV3 clone through CRISPR-Cas9 homology-directed mutagenesis^17^; while the diversity of the library was designed to model decoder-generated sequences of the RSV3 cluster, it also contained fully randomized positions (**Supplementary Fig. 7a**). Screening of the hybridoma antibody library by FACS followed by deep sequencing yielded 19’270 surface-expressed variants of whom 7’470 were confirmed to be antigen-binding (**Supp. Fig. 7b)**. When assessing the probabilities of these novel variants under the VAE model, we found that binding CDRH3s were significantly more likely to be generated than non-binding variants (**Extended Data Fig. 10**). However, since the library also contained a.a. that were not observed in nature, most of its variants were less likely to be generated by our model than naturally occurring sequences (**Extended Data Fig. 10, Supplementary. Figure 8**). Yet, the overlap between the distributions indicated that the VAE model should have been able to generate some of these variants *in silico*. We confirmed this fact by sampling one million latent encodings directly from the respective RSV3 containing cluster of the GMM model. The trained decoder was then used to transform the sampled encodings into distinct position probability matrices from which in turn actual sequences were generated (**Fig. 4a**). This procedure exhaustively sampled the CDRH3 sequence space of RSV3 and yielded 5’005 novel, high quality *in silico* variants that did not occur in the original biological training set. Of these variants, 71 were confirmed by the previous library screen to bind RSV-F, while 25 *in silico* variants were found in the non-binding fraction; resulting in an overall binding accuracy of 74%. Again, the non-binding variants were sampled at a much lower rate than binding variants, indicating that the bulk of the *in silico* generated sequences are likely to be antigen-specific as well (**Fig. 4b, Extended Data Table 2)**.

**Figure 4.**
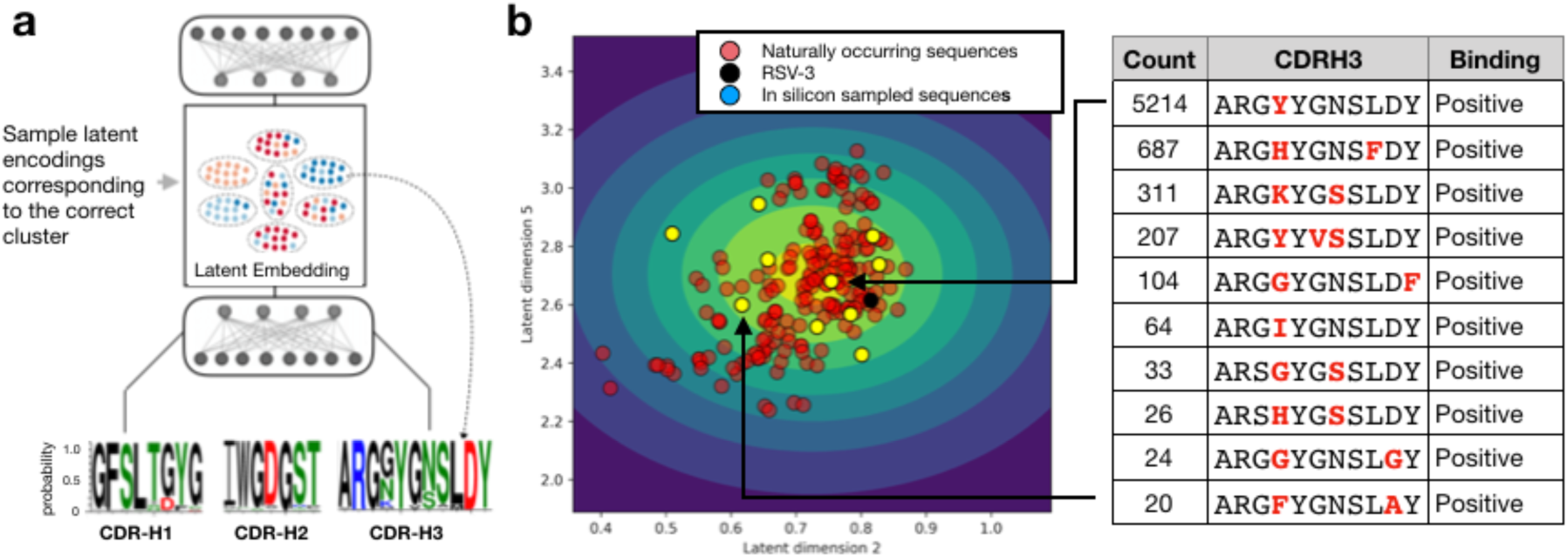
Deep generative modelling and in silico antibody sequence generation. a, Schematic deep generative modeling of antibody sequence space: a cluster is either chosen or randomly sampled and based on the parameters chosen, a random sample is drawn from a multivariate normal distribution. The encoder then translates the encoding into a multivariate multinomial distribution from which a novel sequence is sampled. b, Scatter plot shows the two latent naturally occurring variants, yellow dots show the ten most frequently in-silico sampled encodings that were confirmed to be binding antibodies. The table on the right shows their CDRH3 sequence and its count after 1’000’000 samples. Red letters indicate differences to the initial biological sequence (RSV3, shown in black).

## DISCUSSION

In summary, we show that wide-scale convergence across a range of antigens occurs in the antibody repertoires of mice. Unlike current approaches used to identify clonotypes^18-20^, our VAE approach learns the clustering thresholds based on densities of individual sequence motifs found in the data, thereby forming clusters of varying degrees of similarity. Furthermore, our trained encoding neural network is able to identify these motifs in unseen repertoires in a sensible manner, thereby extending currently existing frameworks for immunodiagnostics^21,22^. Commonly applied methods to detect convergence such as clonotyping based on exact V-gene and J-gene and a CDRH3 similarity threshold (e.g. 80%) are only partly able to recover convergent patterns; in contrast deep learning revealed that convergent antibody sequences utilized various V-genes and J-genes and have dissimilarities of up to >40% in CDRH3. It is therefore likely that the current extent of antigen driven convergence in immune receptor repertoires has been underreported. We are able to discover convergent motifs from sequences buried deep in the repertoire, highlighting again the possibility that as the amount of available sequencing data increases, similar phenomena might be more commonly observed in humans as well. While we focused specifically on the V_H_ region (previous studies have shown them to be sufficient to determine most clonal relationships^23^), we also observed evidence for potential convergence on the V_L_ region; future work using single-cell sequencing^24^ may reveal convergent patterns across the paired V_H_:V_L_ binding domain. We show that our deep generative model allows us to exhaustively sample from the naturally occurring sequence space of antibody repertoires, resulting in the *in silico* generation of antibody variants that retain antigen-binding, a procedure that could be used for engineering antibodies with desired therapeutic development properties^25^. Detection of convergent patterns by deep learning may also enable the discovery of functional and protective antibodies in patients with unique immunological phenotypes (e.g., elite neutralizers of HIV), which could be exploited as immunodiagnostics, therapeutic antibodies or for vaccine immunogen design^21,26-28^.

## METHODS

### Mouse immunizations and RNA isolation from bone marrow

Female BALB/c mice (Charles Rivers) of 6-8 weeks old were separated into cohorts (10-12 mice) based on antigen: hen egg lysozyme (HEL, Sigma Aldrich, 62971-F), ovalbumin (OVA, Hyglos, 321001), blue carrier protein (BCP, Pierce, 771300) and respiratory syncytial virus fusion glycoprotein (RSV-F, expressed internally). For HEL, OVA and BCP, mice were immunized with an initial subcutaneous injection of 200 μg antigen (and 20 μg monophosphoryl lipid A (MPLA, Invivogen, Tlrl-mpla) adjuvant. These mice received zero, one, two or three booster injections for which final immunizations (boost 1, 2 or 3) were done with 50 μg antigen (intraperitoneal injection without any adjuvants). Middle immunizations (boost 1 and/or 2) were done with 50 μg antigen and 20 μg MPLA. For RSV-F, all mice were immunized with 2 booster injections, with each of the three injections consisting of 10 µg for RSV-F and 1% Alum (Thermo Scientific, 77161) adjuvant. For all antigens, sequential injections were interspaced by three weeks. All adjuvants and antigens were prepared and aliquoted before the experiments and mixed in a total volume of 150 μL (for RSV-F: 100µl)) on the days of the corresponding injection. Mice were sacrificed 10 days (for RSV-F: 14 days) after their final immunization and bone marrow was extracted from femurs of hind legs using 2 mM PBS buffer. The isolated bone marrow was then centrifuged at 400 x g at 4 °C for 5 minutes. The supernatant was removed and 1.25 mL of Trizol (Invitrogen, 15596026) was added. The bone marrow was then homogenized using a 18G*2’’ needle (1.2*50 mm). 1 mL of the resulting Trizol solution was then frozen at −80 °C until processing. Mouse cohorts and immunization groups are described in **Supplementary Table 1**. RNA extraction was performed as previously described^12^. Briefly, 1 mL of the homogenate was used as input for the PureLink RNA Mini Kit (Life Technologies, 12183018A). RNA extraction was then conducted according to the manufacturer’s guidelines.

### Antibody repertoire library preparation and deep sequencing

Antibody variable heavy chain (V_H_) libraries for deep sequencing were prepared using a previously established protocol of molecular amplification fingerprinting (MAF), which enables comprehensive error and bias correction^12^. Briefly, a first step of reverse transcription was performed on total RNA using a gene-specific primer corresponding to constant heavy region 1 (CH1) of IgG subtypes and with an overhang region containing a reverse unique molecular identifier (RID). Next, multiplex-PCR is performed on first-strand cDNA using a forward primer set that anneals to framework 1 (FR1) regions of V_H_ and has an overhang region of forward unique molecular identifier (FID) and partial Illumina adapter; reverse primer also contains a partial Illumina sequencing adapter. A final singleplex-PCR step is performed to complete the addition of full Illumina adapters. After library preparation, overall library quality and concentration was determined on a Fragment Analyzer (Agilent). Libraries were then pooled and sequenced on an Illumina MiSeq using the reagent v3 kit (2×300 bp) with 10% PhiX DNA added for quality purposes.

### Data pre-processing and sequence alignment

Before alignment, the raw FASTQ files were processed by a custom CLC Genomics Workbench 10 script. Firstly, low quality nucleotides were removed using the quality trimming option with a quality limit of 0.05. Afterwards, forward and reverse read pairs were merged and resulting amplicons between 350 and 600 base pairs were kept for further analysis. Pre-processed sequences were then error-corrected and aligned using the previously established MAF bioinformatic pipeline ^12^. Downstream analysis was then carried out using Python utilizing both the Tensorflow and scikit-learn libraries.

### Variational autoencoder models of antibody repertoires

Following error and bias correction and alignment of antibody repertoire sequencing data, all discovered combinations of CDRH1, CDRH2 and CDRH3 for each dataset were extracted. In order to process CDRHs of various lengths, sequences were padded with dashes until a certain fixed length (maximum length for each CDRH in the data) was reached. Padded sequences were one-hot encoded, concatenated and used as input into the variational autoencoder (VAE). The VAE model jointly optimizes the ability to reconstruct its input together with a Gaussian mixture model (GMM)-based clustering of the latent space (Fig. 1) according to the following formula:

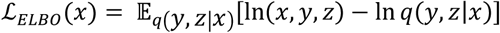

With:

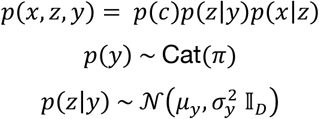

And the following variational approximation of the posterior, where *q*(*z*|*x, y*) is assumed to be distributed according to a gaussian distribution:

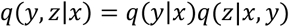

Unlike comparable models^29^, we do not make a mean field approximation when modelling the posterior, which we found to increase model stability. This choice however comes at the expense of considerable increases in computation time, as has been reported before^30^. Specifically, we encode and decode every input sequence as if it would belong to every cluster (indicated through a one-hot encoded cluster label) using shared weights in every layer. The final contributions to the overall loss are then weighted by the separately predicted probabilities *q*(*y*|*x*), which describe the probability of a sequence belonging to a specific cluster (**Extended Data Fig. 1)**. The decoder maps input sequences and concatenated class label into a lower dimensional (d=10) space using two dense layer with rectified linear unit (ReLU) activation followed by the final 10-dimensional layer. Sequences are sampled and recreated from the latent space using the decoder. The decoding network (**Extended Data Fig. 1**) employs two separate dense layers with ReLU activation followed by a dense layer with a linear activation, whose output is reshaped and normalized with a softmax activation in order to reconstruct the probabilities of the initial, one-hot encoded CDRHs. The standard categorical cross-entropy loss is used as the reconstruction term. Every VAE model was trained on a single GPU node of the ETH Zurich parallel computing cluster (*Leonhard)*. Training consisted of 200 epochs for all models using Adam as the optimization algorithm^31^.

### Predicting antigen exposure of single antibody repertoires

Repertoire datasets were split into five folds with each fold being approximately balanced in the number of distinct antigen groups and each dataset appearing only once across all folds. This split was then used to perform a cross-validation procedure in which each of the five folds were set aside as a test set once and the remaining four folds were used as training data. For each of the five training/test splits a separate VAE model was learned by combining all sequences across all repertoires from the training set as input. Clustering assignments or sequences from both the training and the test set were then calculated for the trained model. Based on these cluster labels each repertoire was recoded as an *n*-dimensional vector, where *n* is the number of possible clusters and the *i*-th element indicates the number of sequences mapping to the *i*-th cluster in the given repertoire. These vectors were then used to train and validate linear support vector machines (SVM) in a one-versus-all setting. In order to prevent a more resource-intensive nested cross-validation procedure we decided to not optimize the hyperparameters of the SVMs and instead chose to use the standard values given by scikit-learn’s ‘SVC’ implementation (using a linear kernel). For visualization purposes the results of each cross-validation step were grouped together in one single confusion-matrix (**Fig. 2b**).

### Identification of convergent antigen-associated sequence clusters

In order to identify convergent antigen-associated sequence clusters from antibody repertoires, we performed a non-parametric permutation test in order to determine whether sequence reads were specifically enriched in one cluster given a specific cohort (**Fig. 2c**). In order to account for multiple testing, Bonferroni correction was applied to all p-values in each cohort. We proceeded by randomly choosing one CDRH1-CDRH2-CDRH3 combination and its cognate full-length variable region from each cluster for further validation. Permutation tests were carried out in Python using *mlxtend* and 1000 permutations each.

### Generation of cluster-specific, in-silico variants

Cluster-specific, novel variants were generated in-silico by sampling data points in latent space from a multivariate gaussian distribution, where parameters were given by the respective cluster parameters from the final VAE model. These sampled data points were then fed into the decoding network resulting in position probability matrices for each CDRH (**Fig. 4a**). For each data point a given CDRH1, CDRH2 and CDRH3 was generated. This process was repeated for a million iterations. The log probability of single sequences was approximated by taking the average of 500 samples of the evidence lower bound (ELBO).

### Hybridoma cell culture conditions

All hybridoma cell lines and libraries were cultivated in high-glucose Dulbecco’s Modified Eagle Medium (DMEM; Thermo, 61965026) supplemented with 10% (v/v) heat inactivated fetal bovine serum (FBS; Thermo, 26140079), 100 U/ml penicillin/streptomycin (Pen Strep; Thermo, 15140122), 10 mM HEPES buffer (Thermo, 15630080) and 50 μM 2-Mercaptoethanol (Thermo, 21985023). All hybridoma cultures were maintained in cell culture incubators at a constant temperature of 37C in humidified air with 5% CO_2_. Hybridoma cells were typically cultured in 10 ml of medium in T-25 flasks (TPP, 90026) and passaged every 48/72h. All hybridoma cell lines were confirmed annually to be *Mycoplasma*-free (Universal Mycoplasma Detection Kit, ATCC, 30-1012K). The hybridoma cell line PnP-mRuby/Cas9 was previously published^17^.

### Generation of antibody libraries by CRISPR-Cas9 homology-directed repair

Candidate V_H_ genes were ordered from Twist Bioscience as gene fragments, which were resuspended in 25 μl Tris-EDTA, pH 7.4 (Sigma) prior to use. All oligonucleotides as well as crRNAs and tracrRNA used in this study were purchased from Integrated DNA Technologies (IDT) and adjusted to 100 ſM (oligonucleotides) with Tris-EDTA or to 200 ſM (crRNA/tracrRNAs) with nuclease-free duplex buffer (IDT, 11-01-03-01) prior to use. The homology-directed repair (HDR) donor template used throughout this study was based on the pUC57(Kan)-HEL23-HDR homology donor plasmid, previously described ^16,32^. Importantly, two consecutive stop codons were incorporated into the beginning of the coding regions for the V_H_ and the variable light chain (V_L_) sequences in order to avoid library cloning artefacts and background antibody expression due to unmodified parental vector DNA.

For each candidate V_H_, separate HDR-donor V_L_ libraries were assembled in a stepwise manner by Gibson cloning using the Gibson Assembly Master Mix (NEB, E2611)^33^. When necessary, fragments were amplified using the KAPA Hifi HotStart Ready Mix (KAPA Biosystems, K2602) following manufacturer instructions. First, V_H_ genes were amplified from gene fragments and cloned into the PCR-linearized parental HDR-donor vector (step 1, Extended Data Figure 8). Next, with total bone-marrow RNA of a mouse that was immunized with one of the four respective antigens, reverse transcription was performed using the Maxima Reverse Transcriptase (Thermo, EP0741) with a primer specific for V_L_ constant region. The resulting cDNA was used to amplify the respective V_L_ repertoires in PCR reactions using a degenerate multiplex primer mix, previously described^34^ (**Supplementary Table 6**). V_L_ repertoires were cloned into the PCR-linearized HDR-donor vector created in step 1 for each candidate V_H_ library (step 2) and final libraries were assessed in terms of diversity and background clones. Typically, fixed V_H_ HDR-donor V_L_ libraries had sizes ranging from 30’000 – 80’000 transformants per library.

PnP-mRuby/Cas9 cells were electroporated with the 4D-Nucleofector System (Lonza) using the SF Cell Line 4D-Nucleofector Kit L (Lonza, V4XC-2012) with the program CQ-104. For each HDR-donor library, 10^6^ cells were harvested by centrifugation at 125 x g for 10 min, washed with 1 ml of Opti-MEM Reduced Serum Medium (Thermo, 31985-062) and centrifuged again using the same parameters. The cells were finally resuspended in 100 ſl of nucleofection mix containing 500 pmol of crRNA-J/tracrRNA complex and 20 ſg of HDR-donor plasmid (5.9 kb) diluted in SF buffer. Following electroporation, cells were cultured in 1mL of growth media in 24-well plates (Thermo) for two days and moved to 6-well plates (Costar) containing another 2 mL of growth media for one additional day.

### Screening of hybridoma antibody libraries by flow cytometry

Flow-cytometry analysis and FACS of CRISPR-Cas9 modified hybridomas was performed on a BD LSR Fortessa and BD FACS Aria III (BD Biosciences). Flow cytometry data was analyzed using FlowJo V10 (FlowJo LLC). Three days post-transfection, hybridoma cell libraries specific for one antigen were pooled and enriched for antibody-expressing and antigen-specific cells in consecutive rounds of FACS. Typically, the number of sorted cells from the previous enrichment-step was over-sampled by a factor of 40 in terms of the number of labelled cells for the subsequent sorting-step. For labelling, cells were washed with PBS (Thermo, 10010023), incubated with the labelling antibodies or antigen for 30 min on ice protected from light, washed two times with PBS and analyzed or sorted. The labelling reagents and working concentrations are listed in **Supplementary Table 7**. For cell numbers different from 10^6^, the amount of antibody/antigen as well as the incubation volume were adjusted proportionally. For labelling of RSV-F-specific cells, a two-step labelling procedure was necessary due to the indirect labelling of cells with RSV-F-biotin/Streptavidin-AlexaFluor647.

### Genotyping of monoclonal hybridoma cell lines

Genomic DNA of single-cell sorted and expanded hybridoma clones was isolated from 5×10^5^ cells, which were washed with PBS and resuspended in QuickExtract DNA Extraction Solution (Epicentre, QE09050). Cells were incubated at 68 °C for 15 min and 95 °C for 8 min and the integrated synthetic V_L_-C_k_-2A-V_H_ antibody region was PCR-amplified with flanking primers 5’-CATGTGCCTTTTCAGTGCTTTCTC-3’ and 5’-CTAGATGCCTTTCTCCCTTGACTC-3’ that were specific for the 5’ and 3’ homology arms. From this single amplicon, both V_H_ and V_L_ regions could be Sanger-sequenced using primers 5’-TGACCTTCTCAAGTTGGC-3’ and 5’-GAAAACAACATATGACTCCTGTCTTC-3’, respectively (Microsynth).

### Measurement of antibody specificity by ELISA (cell culture supernatant)

Standard sandwich enzyme-linked immunoabsorbance assays (ELISAs) were performed to measure the specificity of hybridoma cell line supernatants containing secreted IgG. High binding 96-well plates (Costar, CLS3590) were coated over night with 4 ug/ml of antigen in PBS at 4C. The plates were then blocked for two hours at room temperature with PBS containing 2 % (m/v) non-fat dried milk powder (AppliChem, A0830) and 0.05 % (v/v) Tween-20 (AppliChem, A1389). After blocking, plates were washed three times with PBS containing 0.05 % (v/v) Tween-20 (PBST). Cell culture supernatants were 0.2 μm sterile-filtrated (Sartorius, 16534) and serially diluted across the plate (1:3 steps) in PBS supplemented with 2 % (m/v) milk (PBSM), starting with non-diluted supernatants as the highest concentrations. Plates were incubated for one hour at room temperature and washed three times with PBST. HRP-conjugated rat monoclonal [187.1] anti-mouse kappa light chain antibody (abcam ab99617) was used as secondary detection antibody, concentrated at 0.7 μg/ml (1:1500 dilution from stock) in PBSM. Plates were incubated at room temperature for one hour again, followed by three washing steps with PBST. ELISA detection was performed using the 1-Step Ultra TMB-ELISA Substrate Solution (Thermo, 34028) and reaction was terminated with 1 M H_2_SO_4_. Absorption at 450 nm was measured with the Infinite 200 PRO NanoQuant (Tecan) and data were analyzed using Prism V8 (Graphpad).

### Measurement of antibody serum titers by ELISA

Serum titers were measured in a similar manner as described above with the following exceptions: (1) Plates were coated with either 10μg/mL (OVA, BCP, HEL) or 2μg/mL of antigen (RSV-F) dissolved in PBS. (2) OVA, BCP and HEL serum ELISAs were blocked with 300μL/well of PBS with 3% BSA, 3%FBS, 0.05% Tween and 0.05% Proclin and were incubated overnight at 4 °C.

### RSV3 CDRH3 library generation

RSV3 CDRH3 libraries were generated following CRISPR-Cas9 homology-directed mutagenesis, as previously described ^17^. Briefly, a single-stranded oligonucleotide (ssODN) encoding a nucleotide sequence that put the endogenous CDRH3 out of frame and contained a highly efficient CRISPR targeting sequence was incorporated into the genomic CDRH3 locus of the RSV3 cell line by CRISPR/Cas9-mediated HDR, using reagents crRNA DN_RSV3_H3-3 and 500 pmol of DN_RSV3-OOF ssODN HDR-template (RSV3-OOF cell line, **Supplementary Fig. 7a, Supplementary Table 7**). Next, 4 × 10^6^ RSV3-OOF cells were transfected with crRNA-DM1 and 500pmol of ssODN encoding a CDRH3 library template per 1 × 10^6^ cells to generate the RSV3 CDRH3 in silico library. Transfected cells were subsequently sorted in two consecutive steps for antibody expression and specificity/negativity towards RSV-F (as described above) to enrich for a pure RSV-F-specific cell population and an RSV-F-negative cell population.

### Deep sequencing of RSV3 CDRH3 libraries

Sample preparation for deep sequencing was performed following a previously established two-step primer extension protocol [23]. Genomic DNA was extracted from 5 × 10^6^ cells of IgG+, IgG+/RSVF+ and IgG+/RSVF-enriched cell populations using the PureLink^TM^ Genomic DNA Mini Kit (Thermo, K182001). Extracted genomic DNA was amplified in a first PCR using a forward primer binding to the beginning of FR3 and a reverse primer binding to the intronic region located ∼40 bp downstream of the end of the J-gene.

PCRs were performed with Q5 High-Fidelity DNA polymerase (NEB, M0491L) in 8 parallel 50ul reactions with the following cycle conditions: 98 °C for 30 s; 20 cycles of 98 °C for 10 s, 64 °C for 20 s, 72 °C for 20 s; final extension 72 °C for 2 min; 4 °C storage. PCR products were subsequently concentrated using the DNA Clean and Concentrator Kit (Zymo, D4013) followed by 1.2X SPRIselect (Beckman Coulter, B22318) left-sided size selection. Total PCR1 product (∼700 ng) was amplified in a second PCR step, which added extension-specific full-length Illumina adapter sequences to each library by choosing 3 different index reverse primers (DM142-144, using DM125 as the forward primer). Cycle conditions were as follows: 98 °C for 30 s; 2 cycles of 98 °C for 10 s, 40 °C for 20 s, 72 °C for 1 min; 15 cycles of 98 °C for 10 s, 65 °C for 20 s, 72 °C for 30 s; final extension 72 °C for 2 min; 4 °C storage. PCR2 products were subsequently concentrated again using the DNA Clean and Concentrator Kit and run on a 1% agarose gel. Product bands of the correct size (∼400 bp) were gel-purified using the Zymoclean™ Gel DNA Recovery Kit (Zymo, D4008) and subsequently analyzed by capillary electrophoresis (Fragment analyzer, Agilent). After quantitation, libraries were pooled accordingly and sequenced on a MiSeq System (Illumina) with the paired-end 2×300bp kit.

## AUTHOR CONTRIBUTIONS

S.F., D.N., T.A.K., D.M. and S.T.R. designed experiments. T.A.K, A.R.G.D.V., L.C. performed mouse-related experiments. S.F., A.R.G.D.V. and L.C. prepared sequencing libraries. D.N., C.P. conducted preliminary experiments. D.N, L.E. conducted experimental antibody library screening. S.F. was responsible for the bioinformatics pipeline. S.F. analyzed deep sequencing data and developed machine learning and deep learning models. S.F., D.N. prepared figures. S.F., D.N., and S.T.R. wrote the paper.

### ACKNOWLEDGMENTS

We acknowledge the ETH Zurich D-BSSE Single Cell Unit and the Genomics Facility Basel for excellent support and assistance. This work was supported by the European Research Council Starting Grant 679403 (to S.T.R.) and ETH Zurich Research Grants (to S.T.R.) and NCCR Molecular Systems Engineering (to S.T.R). The professorship of S.T.R. is supported by an endowment from the S. Leslie Misrock Foundation.

## COMPETING INTERESTS

ETH Zurich has filed for patent protection on the technology described herein, and S.F. and S.T.R. are named as co-inventors on this patent (United States Patent and Trademark Office Provisional Application: 62/843,010).

## DATA AVAILABILITY

The raw FASTQ files from deep sequencing that support the findings of this study will be deposited (following peer-review and publication) in the Sequence Read Archive (SRA) with the primary accession code(s) <code(s) (https://www.ncbi.nlm.nih.gov/sra)>. Additional data that support the findings of this study are available from the corresponding author upon reasonable request.

## CODE AVAILABILITY

Deep learning models were built in Python v3.6.4 using TensorFlow v1.12.0. Models will be available on a github repository (following peer-review and publication).

## REFERENCES

1 Wang, B. et al. Functional interrogation and mining of natively paired human VH:VL antibody repertoires. Nature biotechnology 36, 152–155, doi:10.1038/nbt.4052 (2018).

2 Briney, B., Inderbitzin, A., Joyce, C. & Burton, D. R. Commonality despite exceptional diversity in the baseline human antibody repertoire. Nature 566, 393–397, doi:10.1038/s41586-019-0879-y (2019).

3 Greiff, V. et al. Systems Analysis Reveals High Genetic and Antigen-Driven Predetermination of Antibody Repertoires throughout B Cell Development. Cell Rep 19, 1467–1478, doi:10.1016/j.celrep.2017.04.054 (2017).

4 Soto, C. et al. High frequency of shared clonotypes in human B cell receptor repertoires. Nature 566, 398–402, doi:10.1038/s41586-019-0934-8 (2019).

5 Croote, D., Darmanis, S., Nadeau, K. C. & Quake, S. R. High-affinity allergen-specific human antibodies cloned from single IgE B cell transcriptomes. Science 362, 1306–1309, doi:10.1126/science.aau2599 (2018).

6 Setliff, I. et al. Multi-Donor Longitudinal Antibody Repertoire Sequencing Reveals the Existence of Public Antibody Clonotypes in HIV-1 Infection. Cell host & microbe 23, 845–854 e846, doi:10.1016/j.chom.2018.05.001 (2018).

7 Parameswaran, P. et al. Convergent antibody signatures in human dengue. Cell host & microbe 13, 691–700, doi:10.1016/j.chom.2013.05.008 (2013).

8 Truck, J. et al. Identification of antigen-specific B cell receptor sequences using public repertoire analysis. Journal of immunology 194, 252–261, doi:10.4049/jimmunol.1401405 (2015).

9 Glanville, J. et al. Identifying specificity groups in the T cell receptor repertoire. Nature 547, 94–98, doi:10.1038/nature22976 (2017).

10 Dash, P. et al. Quantifiable predictive features define epitope-specific T cell receptor repertoires. Nature 547, 89 (2017).

11 Friedensohn, S., Khan, T. A. & Reddy, S. T. Advanced Methodologies in High-Throughput Sequencing of Immune Repertoires. Trends in Biotechnology 35, 203–214, doi:10.1016/j.tibtech.2016.09.010 (2017).

12 Khan, T. A. et al. Accurate and predictive antibody repertoire profiling by molecular amplification fingerprinting. Sci Adv 2, e1501371, doi:10.1126/sciadv.1501371 (2016).

13 Davidsen, K. et al. Deep generative models for T cell receptor protein sequences. Elife 8, doi:10.7554/eLife.46935 (2019).

14 Riesselman, A. J., Ingraham, J. B. & Marks, D. S. Deep generative models of genetic variation capture the effects of mutations. Nature methods 15, 816–822, doi:10.1038/s41592-018-0138-4 (2018).

15 Sinai, S., Kelsic, E., Church, G. M. & Nowak, M. A. Variational auto-encoding of protein sequences. arXiv preprint 1712.03346 (2017).

16 Pogson, M., Parola, C., Kelton, W. J., Heuberger, P. & Reddy, S. T. Immunogenomic engineering of a plug-and-(dis)play hybridoma platform. Nature communications 7, 12535, doi:10.1038/ncomms12535 (2016).

17 Mason, D. M. et al. High-throughput antibody engineering in mammalian cells by CRISPR/Cas9-mediated homology-directed mutagenesis. Nucleic acids research 46, 7436–7449, doi:10.1093/nar/gky550 (2018).

18 Gupta, N. T. et al. Hierarchical Clustering Can Identify B Cell Clones with High Confidence in Ig Repertoire Sequencing Data. Journal of immunology 198, 2489–2499, doi:10.4049/jimmunol.1601850 (2017).

19 Glanville, J. et al. Naive antibody gene-segment frequencies are heritable and unaltered by chronic lymphocyte ablation. Proceedings of the National Academy of Sciences of the United States of America 108, 20066–20071, doi:10.1073/pnas.1107498108 (2011).

20 Greiff, V., Miho, E., Menzel, U. & Reddy, S. T. Bioinformatic and Statistical Analysis of Adaptive Immune Repertoires. Trends in immunology 36, 738–749, doi:10.1016/j.it.2015.09.006 (2015).

21 Greiff, V. et al. A bioinformatic framework for immune repertoire diversity profiling enables detection of immunological status. Genome Med 7, 49, doi:10.1186/s13073-015-0169-8 (2015).

22 Emerson, R. O. et al. Immunosequencing identifies signatures of cytomegalovirus exposure history and HLA-mediated effects on the T cell repertoire. Nat Genet 49, 659–665, doi:10.1038/ng.3822 (2017).

23 Zhou, J. Q. & Kleinstein, S. H. Cutting edge: ig H chains are sufficient to determine most B cell clonal relationships. The Journal of Immunology 203, 1687–1692 (2019).

24 Laserson, U. et al. High-resolution antibody dynamics of vaccine-induced immune responses. Proceedings of the National Academy of Sciences of the United States of America 111, 4928–4933, doi:10.1073/pnas.1323862111 (2014).

25 Jain, T. et al. Biophysical properties of the clinical-stage antibody landscape. Proceedings of the National Academy of Sciences of the United States of America 114, 944–949, doi:10.1073/pnas.1616408114 (2017).

26 Jardine, J. G. et al. HIV-1 broadly neutralizing antibody precursor B cells revealed by germline-targeting immunogen. Science 351, 1458–1463, doi:10.1126/science.aad9195 (2016).

27 Sesterhenn, F. et al. Boosting subdominant neutralizing antibody responses with a computationally designed epitope-focused immunogen. PLoS Biol 17, e3000164, doi:10.1371/journal.pbio.3000164 (2019).

28 Horns, F., Dekker, C. L. & Quake, S. R. Memory B Cell Activation, Broad Anti-influenza Antibodies, and Bystander Activation Revealed by Single-Cell Transcriptomics. Cell Reports 30, 905–913. e906 (2020).

29 Jiang, Z., Zheng, Y., Tan, H., Tang, B. & Zhou, H. Variational Deep Embedding: An Unsupervised and Generative Approach to Clustering.

30 Kingma, D. P., Mohamed, S., Rezende, D. J. & Welling, M. In Advances in neural information processing systems. 3581–3589.

31 Kingma, D. P. & Ba, J. Adam: A method for stochastic optimization. arXiv preprint 1412.6980 (2014).

32 Parola, C. et al. Antibody discovery and engineering by enhanced CRISPR-Cas9 integration of variable gene cassette libraries in mammalian cells. MAbs 11, 1367–1380, doi:10.1080/19420862.2019.1662691 (2019).

33 Gibson, D. G. et al. Enzymatic assembly of DNA molecules up to several hundred kilobases. Nature methods 6, 343–U341, doi:10.1038/Nmeth.1318 (2009).

34 Reddy, S. T. et al. Monoclonal antibodies isolated without screening by analyzing the variable-gene repertoire of plasma cells. Nature biotechnology 28, 965–969, doi:10.1038/nbt.1673 (2010).

35 Lefranc, M. P. Nomenclature of the human immunoglobulin heavy (IGH) genes. Exp Clin Immunogenet 18, 100–116, doi:49189 (2001).

