## Supplementary Material for "Convergent selection in antibody repertoires is revealed by deep learning"

**EXTENDED DATA FIGURES and TABLES**

**SUPPLEMENTARY FIGURES AND TABLES**

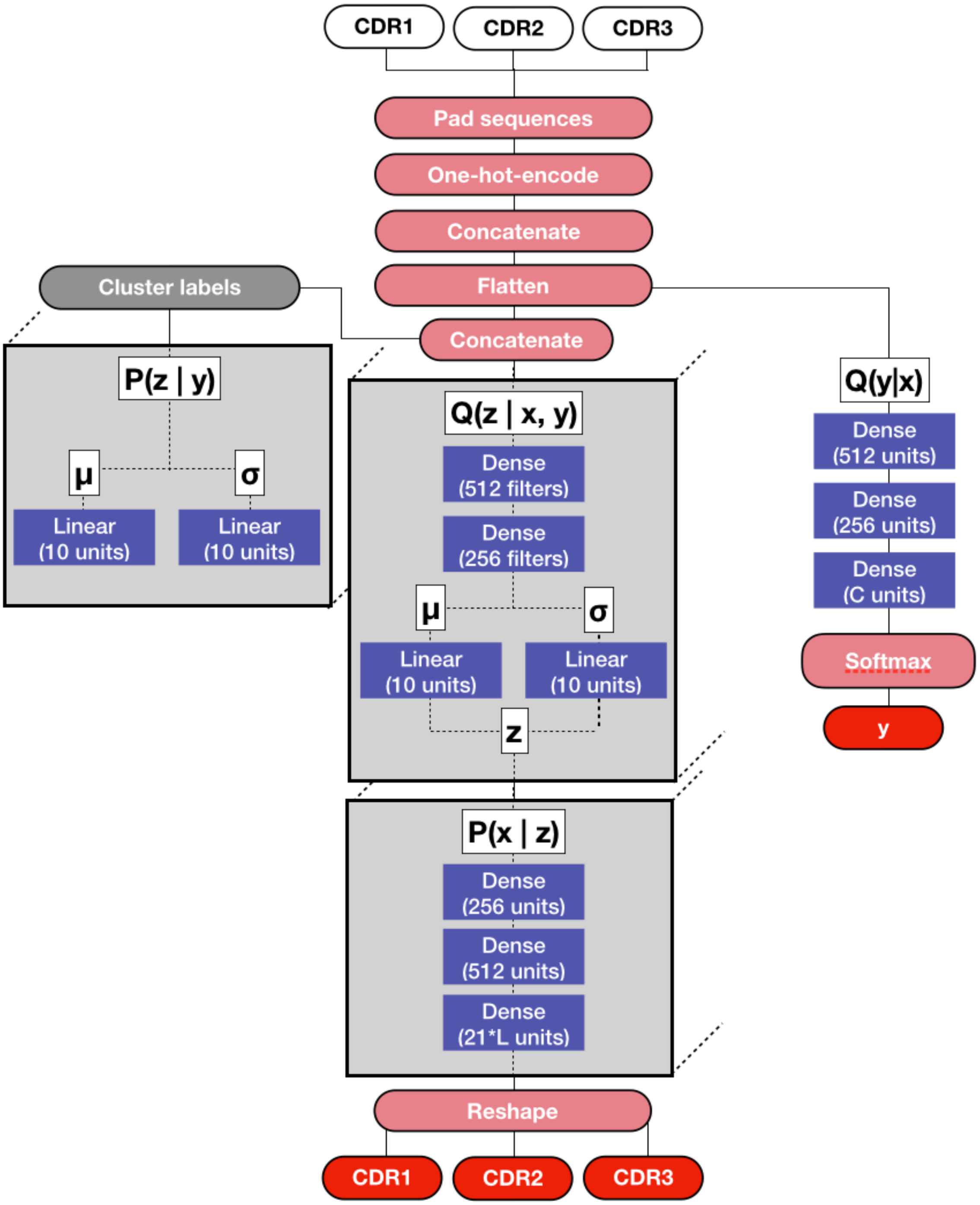


**Extended Data Figure 1. Deep neural network architecture of variational autoencoder.** Grey boxes indicate the input into the model, while red boxes indicate various (mathematical) operations. Purple boxes highlight the trainable layers of the model. Dark red indicates the output of the model. Grey boxes contain layers whose weights are shared across all cluster dimensions.

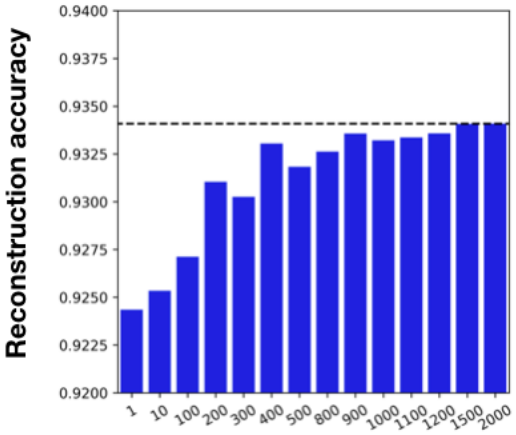


**Extended Data Figure 2. Reconstruction accuracy of variational autoencoder.**
Bar plots show the achieved reconstruction accuracy as a function of the number of clusters. Increasing the of amount clusters (k) results in an increase in the reconstruction accuracy, with diminishing returns after k = 2000.

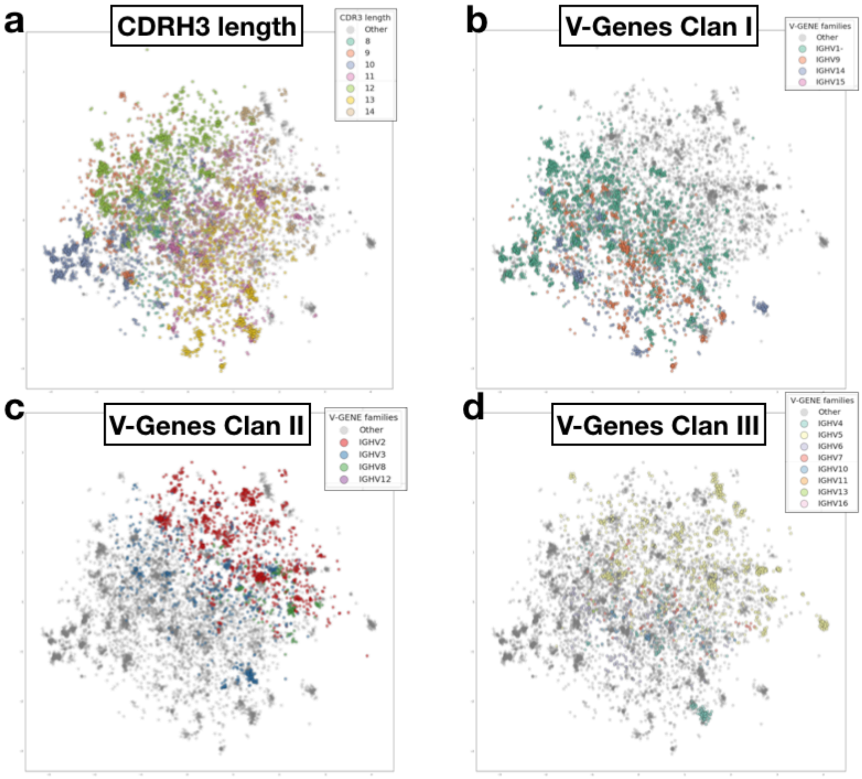


**Extended Data Figure 3. Latent space captures sequence features of antibody repertoires.**
Ten-dimensional latent space of two antibody repertoires (OVA (2C) and RSV-F (2C), shown in **Fig. 2a**) visualized by principal component analysis (PCA). a, Colored dots indicate sequence length**; b-d,** or germline V-gene family membership stratified according to clans based on Immunogenetics Database[35].

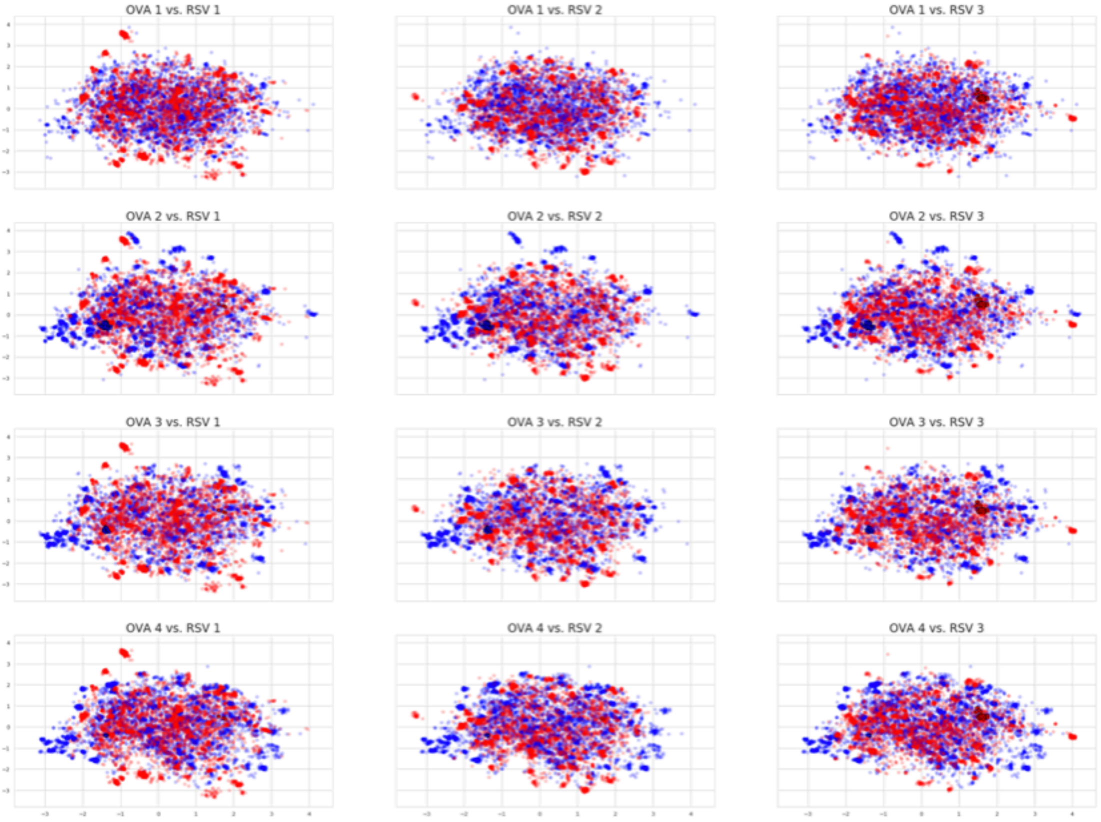


**Extended Data Figure 4. Additional latent space visualizations of antibody repertoires.**
PCA visualization of ten-dimensional latent space of antibody repertoires comparisons from different mice. Blue and red dots indicate sequences belonging to OVA or RSV-F repertoires, respectively.

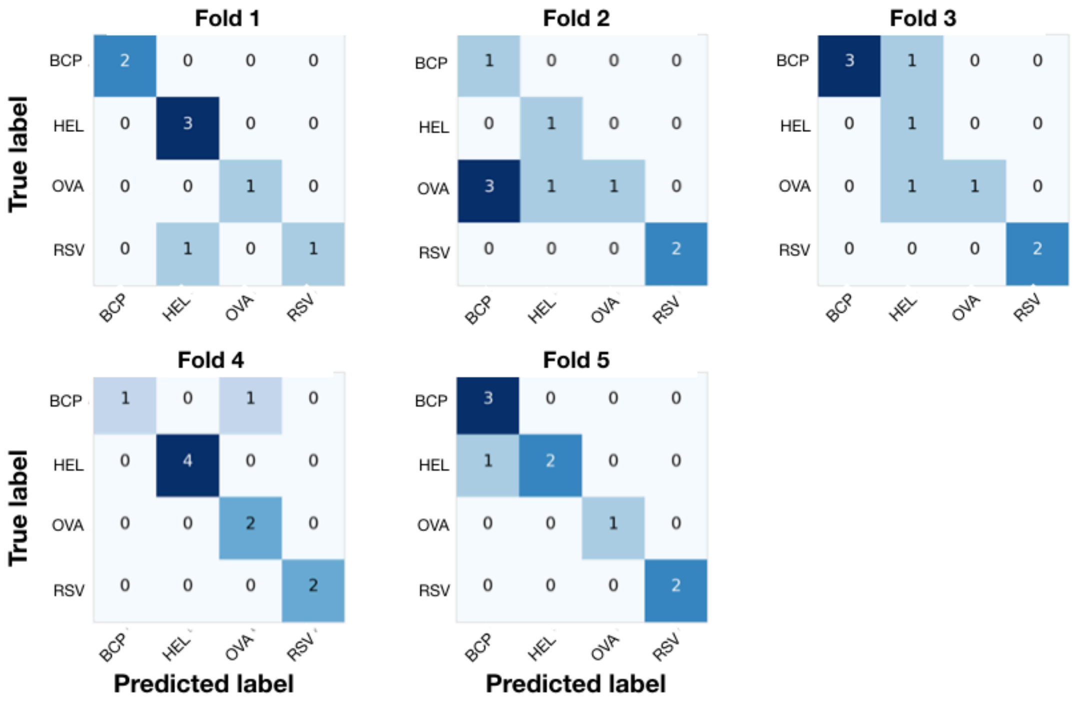


**Extended Data Figure 5. VAE convergent cluster-based prediction of antigen exposure.**
Confusion matrices show the prediction results of each VAE model for each of the 5-folds using aggregated convergent cluster labels as features. Folds are produced in such a manner that no dataset gets evaluated twice.

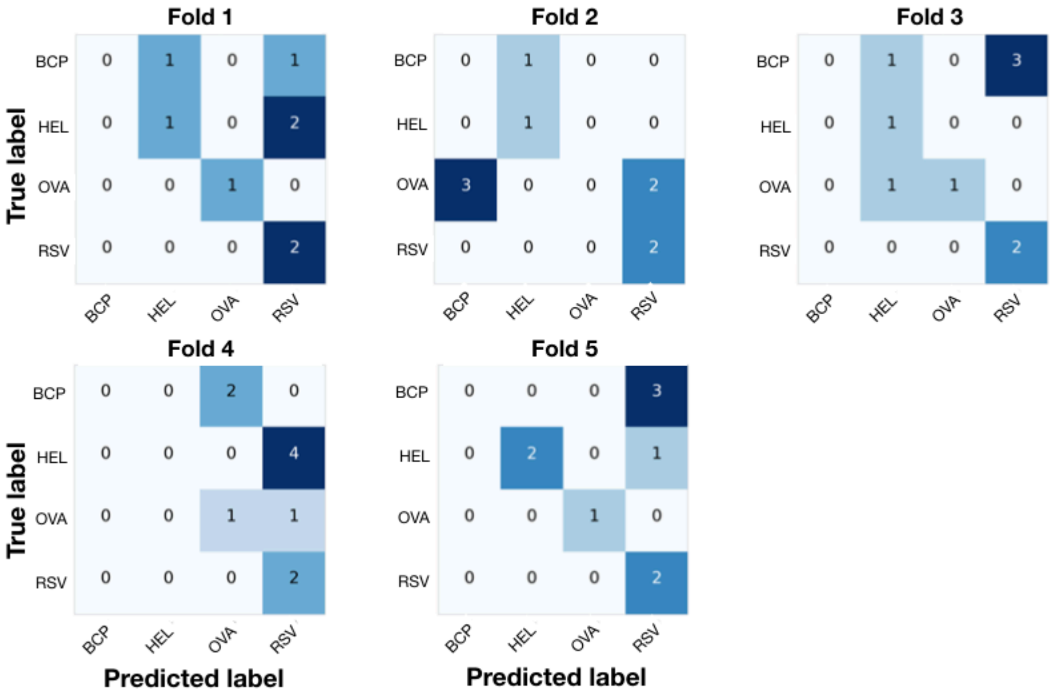


**Extended Data Figure 6. Public clone-based prediction of antigen exposure.**
Confusion matrices show the prediction results of each model for each of the 5-folds using public clone sequences as features. Folds are produced in such a manner that no dataset gets evaluated twice.

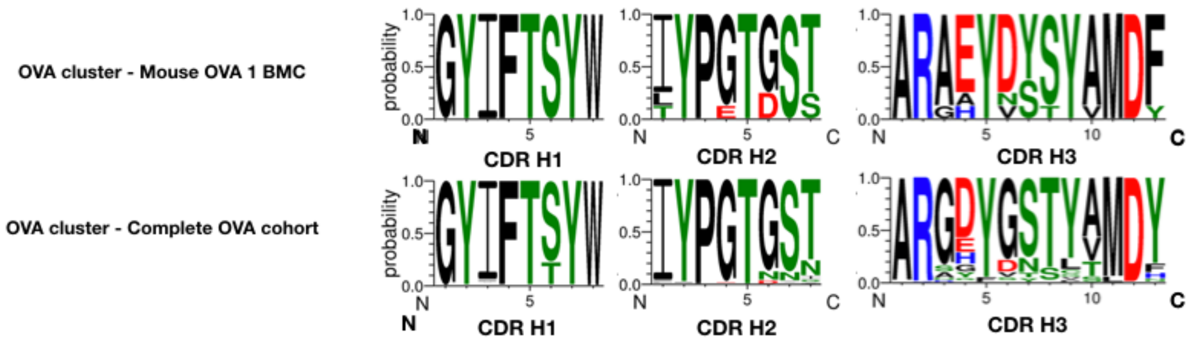


**Extended Data Figure 7. Exemplary convergent clusters containing sequences from single or multiple mouse repertoires.**

Motifs highlight convergent sequences found in one exemplary OVA-cluster from one mouse (top) or all mice from OVA cohort (bottom).

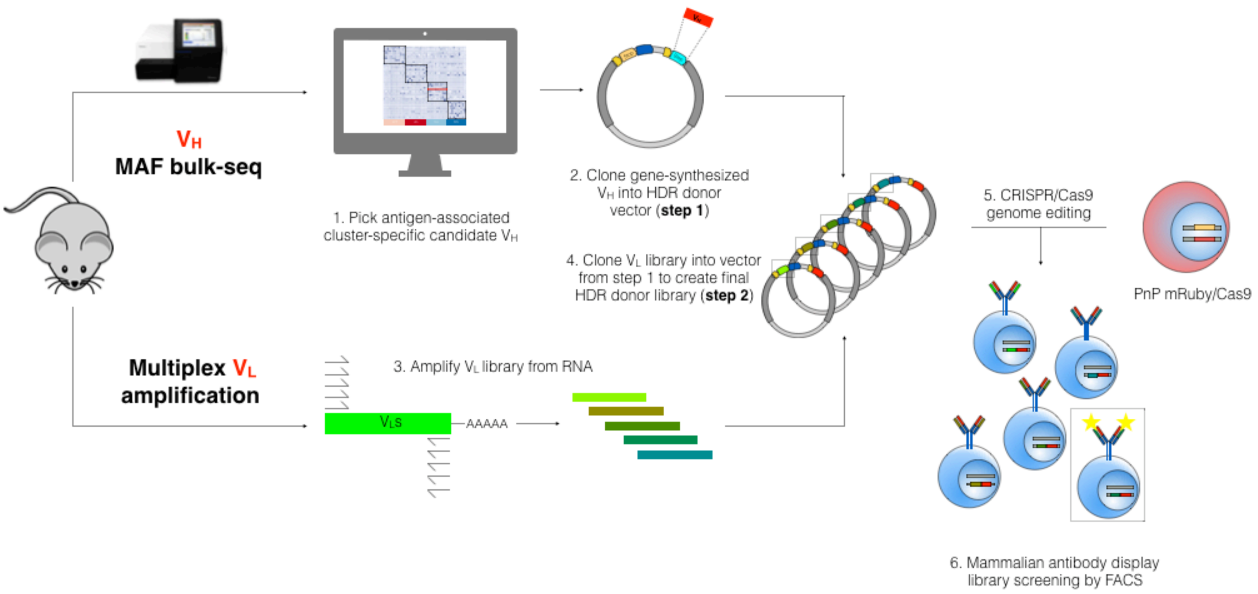


**Extended Data Figure 8. Hybridoma antibody library screening workflow.**

Candidate convergent $V_H$ chains are selected for each antigen. Sequences are gene-synthesized and cloned into the homology-directed repair (HDR) donor vector (step 1). For each antigen, the $V_L$ chain repertoire is amplified from the RNA of a mouse that was immunized with the same antigen by multiplex PCR. The resulting $V_L$ library is then cloned into the HDR donor vector from step 1 (step 2). The resulting HDR donor libraries are then used for CRISPR-Cas9-based integration into the PnP mRuby/Cas9 cells (as described in [16]), thereby resulting in a library of hybridoma cells that express antibodies with candidate convergent $V_H$ chains and diverse $V_L$ library. Hybridoma antibody library is screened for antigen-specific binding variants and selected by FACS.

| Antigen | V-Gene | J-Gene | CDR1 | CDR2 | CDR3 | Public Clone | Convergent GMM-VAE | Binder |
|---|---|---|---|---|---|---|---|---|
| **OVA1** | IGHV1-80 | IGHJ3 | GYAFSSYW | INPGDGDT | ARGGYGSFAY | No | Yes(146) | Yes |
| **OVA2** | IGHV1-76 | IGHJ4 | GYIFTSYW | IYPGTGSS | ARAEYDSSYAMDF | No | Yes(98) | Yes |
| **OVA3** | IGHV1-76 | IGHJ2 | GYIFTSYW | IYPGTGST | ARGDYGSSYFDY | Yes | Yes(142) | Yes |
| **OVA4** | IGHV1-76 | IGHJ4 | GYIFTSYW | IYPGTGST | ARNGDYAMDY | No | Yes(150) | Yes |
| **OVA5** | IGHV1-53 | IGHJ3 | GYTFTSYY | INPSNGGT | TRGGYGYGSFAY | No | Yes(119) | Yes |
| **OVA6** | IGHV1-76 | IGHJ4 | GYIFTSYW | IYPGTGST | AITTVNAMDY | No | Yes(150) | Yes |
| **RSV-F1** | IGHV14-4 | IGHJ4 | GFNIKDYY | IDPENGDT | NAWDGNLDY | No | Yes(136) | Yes |
| **RSV-F2** | IGHV8-12 | IGHJ2 | GFSLSTSGMG | IYWDDDK | ARRSYGSSLDY | Yes | Yes(60) | Yes |
| **RSV-F3** | IGHV2-6-7 | IGHJ2 | GFSLTDYG | IWGDGST | ARGNYGNSLDY | No | Yes(199) | Yes |
| **RSV-F4** | IGHV9-2-1 | IGHJ3 | GYTFTDYS | INTETGEP | ARGGNPAWFAY | No | Yes(98) | NA |

**Extended Data Table 1. Convergent antibody sequences screened for antigen-binding.**
Table shows convergent sequences experimentally screened for antigen-binding. The three rightmost columns indicate whether a sequence could have been identified by the respective method. A sequence would have been discovered as public clone if it is shared with at least one other mouse in its cohort, but was not observed in any other antigen cohort. Number in parentheses indicates the number of sequences found in the convergent cluster.

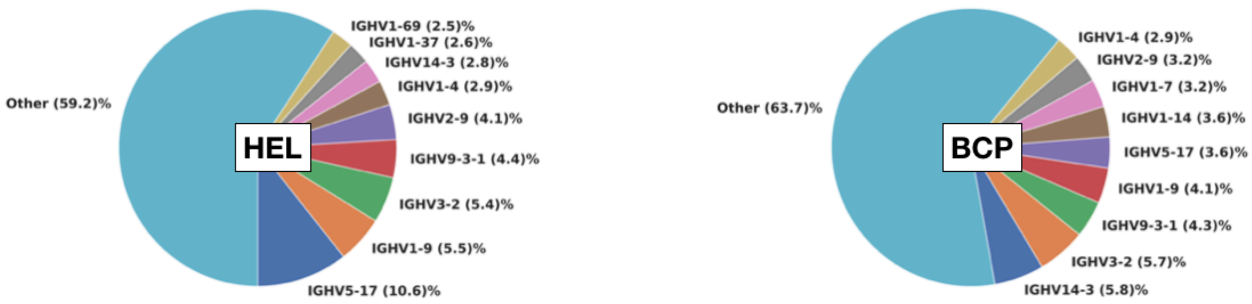


**Extended Data Figure 9. Variable germline (V-gene) usage of convergent antibody sequences.**
Pie charts show the nine most utilized V-gene germlines in convergent clones for HEL and BCP cohorts.

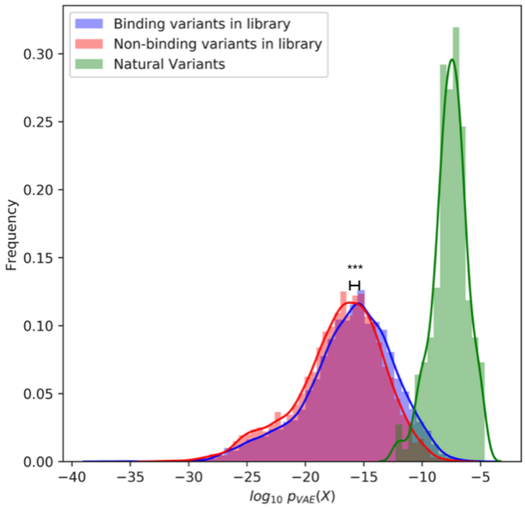


**Extended Data Figure 10. Sampling results from RSV3 generated CDRH3 library.**
Histograms show how likely sequences from the positive (blue) and negative (red) fraction of the RSV3 CDRH3 library screen are to occur according to the VAE decoder model. Positive variants are slightly but significantly (P <0.001, Mann-WhitneyU test) more likely to occur. The green histogram to the right depicts the probabilities of the variants observed in the biological repertoires.

| Count | CDR-H3 | Binding |
|---|---|---|
| 4788 | CARGYYGSSLDYW | Unkown |
| 4382 | CARGGYGNSLDYW | Unkown |
| 2947 | CARGGYGSSLDYW | Unkown |
| 1905 | CARGYYGNSFDYW | Unkown |
| 1750 | CARGNYGNSLDYW | Unkown |
| 1529 | CARGFYGNSLDYW | Unkown |
| 1452 | CARGNYGSSLDYW | Unkown |
| 1395 | CARGHYGNSLDYW | Unkown |
| 1390 | CARGYYGSSFDYW | Unkown |
| 1327 | CARGGYGNSFDYW | Unkown |
| 68 | CAREYYGSSLDYW | Negative |
| 43 | CARDFYGSSLDYW | Negative |
| 21 | CARRYYGSSFDYW | Negative |
| 21 | CARGVYGSSLDYW | Negative |
| 16 | CARRNYGNSLDYW | Negative |
| 11 | CARRNYGNSLDYW | Negative |
| 9 | CARRHYGSSLDYW | Negative |
| 7 | CARRGYGSSFDYW | Negative |
| 3 | CARGVYVSSFDYW | Negative |
| 3 | CARYFYGNSLDYW | Negative |

**Extended Data Table 2. Overview of sampled sequences of RSV3 cluster in silico generation.**
Table shows the top ten most commonly sampled CDRH3 sequences of RSV3 generated library, which are either unknown to bind (blue) or proven to not bind RSV-F (red), and their counts after 1'000'000 samples.

### SUPPLEMENTARY FIGURES AND TABLES

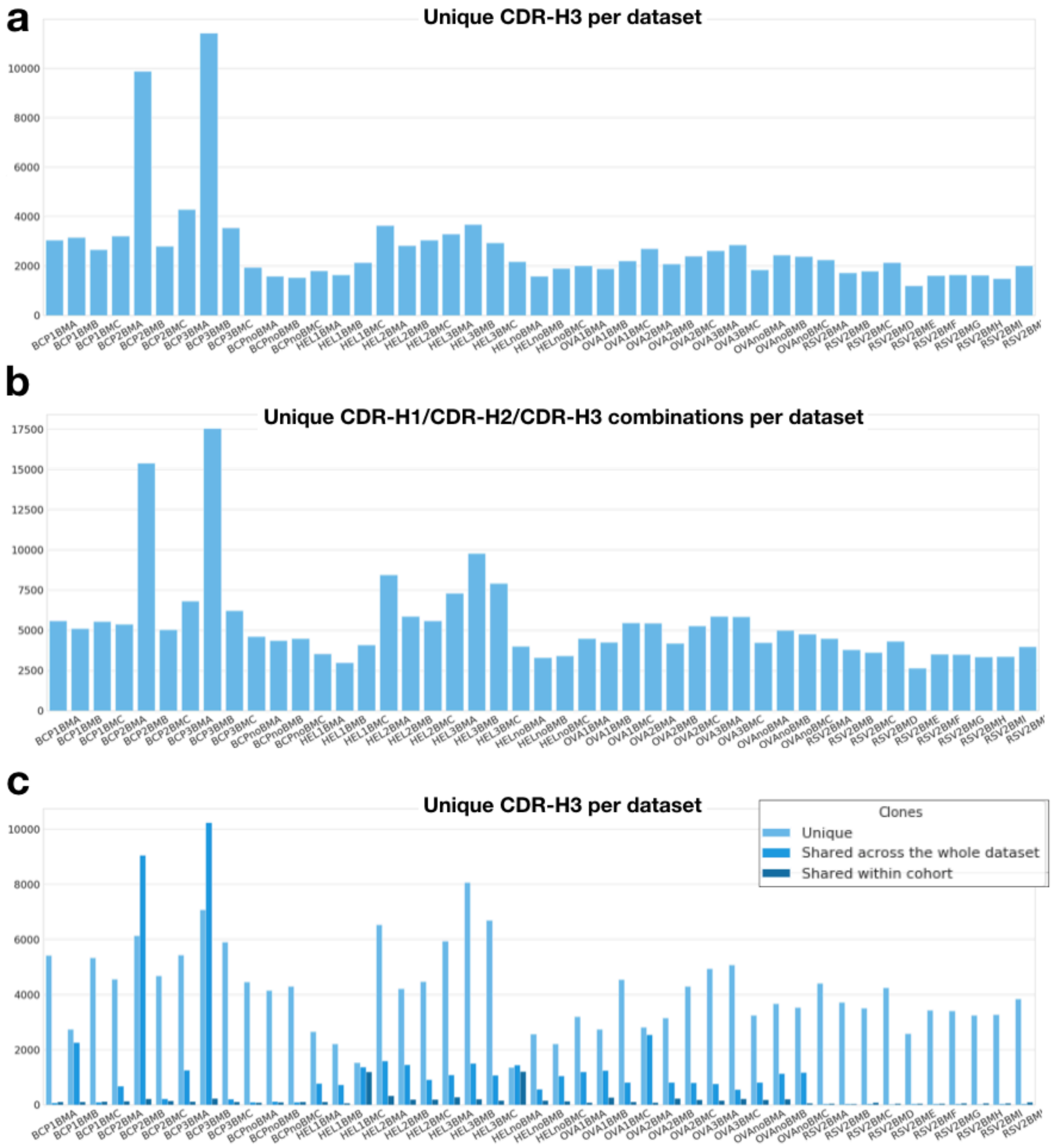


**Supplementary Figure 1. Deep sequencing results of antibody repertoires.**
a, Bar plot shows the number of unique CDRH3 sequences found in each dataset after pre-processing and alignment. b, Bar plot shows the number of unique combinations of all three CDRHs found in each dataset. c, Bar plots show the number of private (sequences only found in the given repertoire) and shared (public) CDRH3 combinations (found in other repertoires).

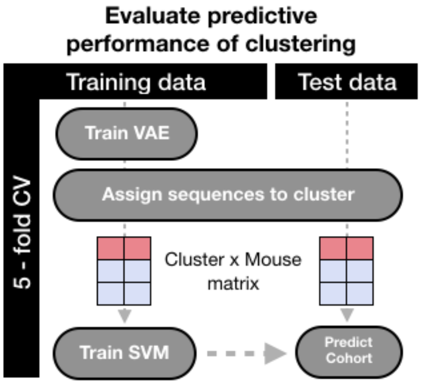


**Supplementary Figure 2. Cross-validation procedure for support vector machine prediction of antigen exposure.**
Antibody repertoires are transformed into vectors containing the enrichment of the learned sequence clusters in latent space. Recoded vectors are used as input for a linear support vector machine (SVM).

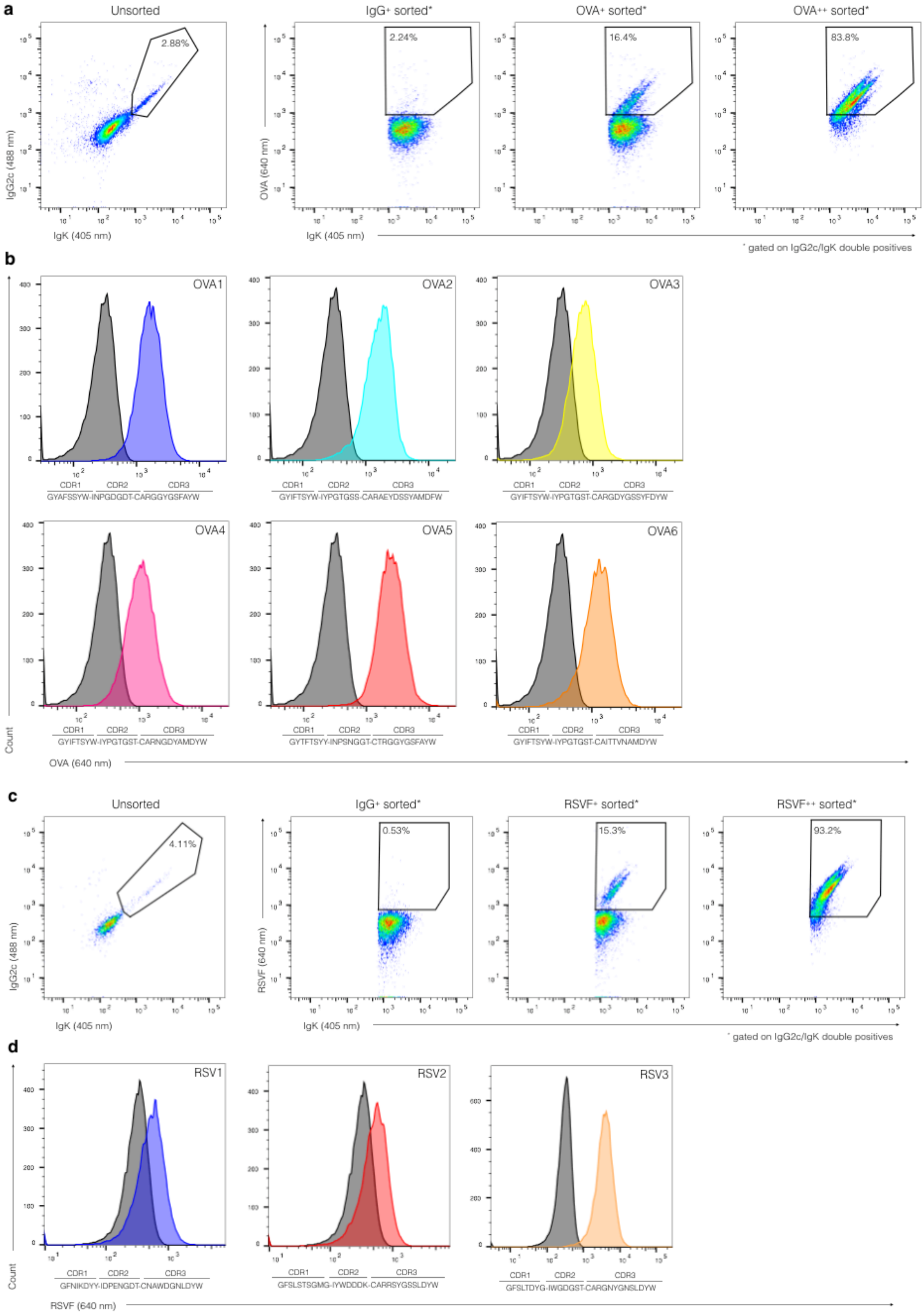


**Supplementary Figure 3. Flow cytometry analysis of hybridoma libraries.**
Flow cytometry analysis of hybridoma cell libraries for a-b, OVA and c, d RSV-F. Sequential library enrichment dot plots are shown in a, c. Respective antigen-specific monoclonal cell lines are shown in histogram plots b, d with respect to a negative control cell line that is not-specific for the given antigen.

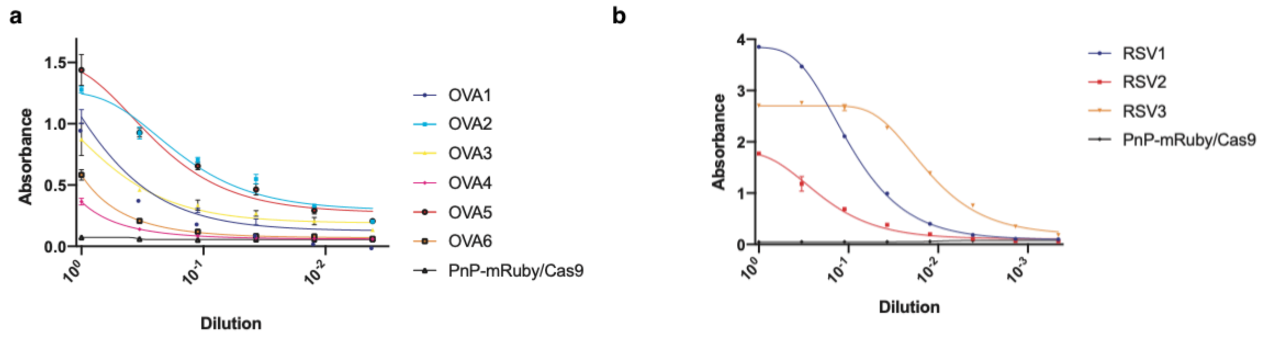


**Supplementary Figure 4. ELISA data of convergent sequences confirmed to be antigen-specific.** Supernatant ELISA profiles of antigen-specific hybridoma monoclonal cell lines are shown for a, OVA b, and RSV-F. Starting cell line PnP-mRuby/Cas9 was used as negative control.

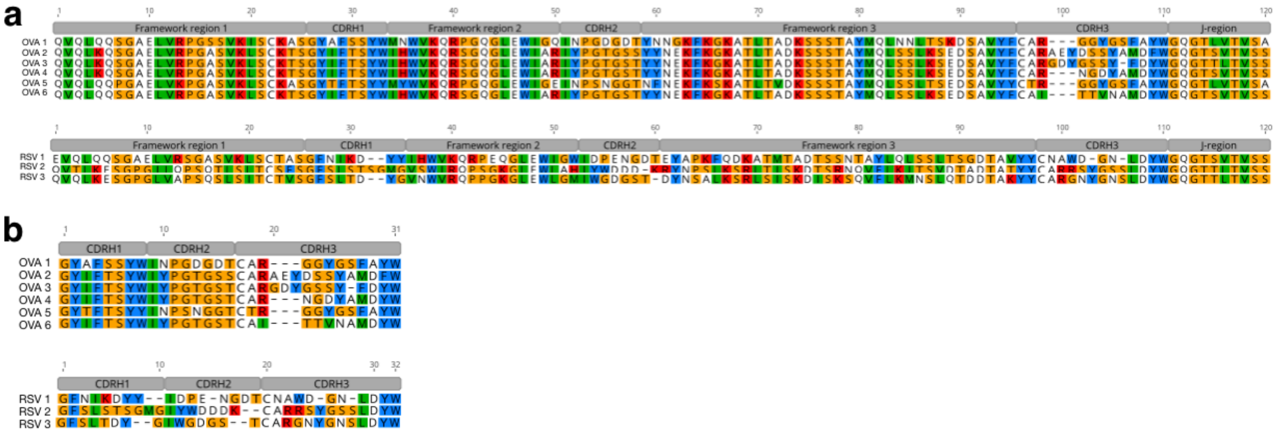


**Supplementary Figure 5. Alignments of convergent sequences confirmed to be antigen-specific.** $V_H$ amino acid alignments for antigen-specific antibodies. a, Full-length VDJ-alignments are shown for OVA and RSV variants. b, Concatenated CDRH1-CDRH2-CDRH3 amino acid alignments for OVA and RSV-F are shown. Color-code used is derived from the Clustal coloring scheme with software Geneious V 10.2.6.

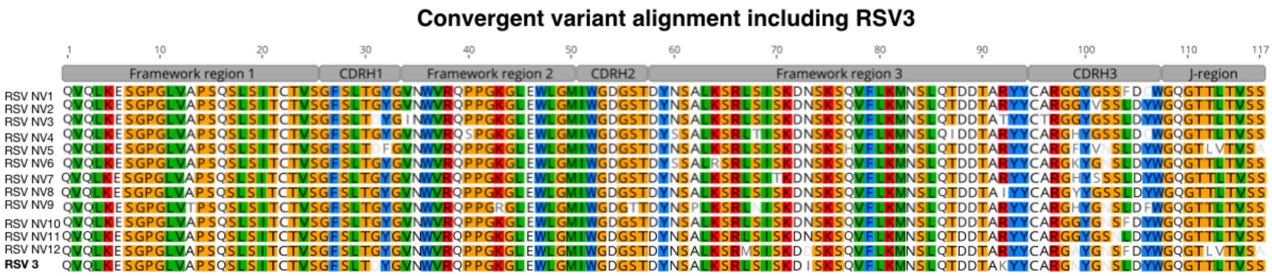


**Supplementary Figure 6. Alignment of convergent variants from RSV3 cluster.** $V_H$ amino acid alignments for convergent natural variants (NV) from RSV3 cluster. Color-code used is derived from the Clustal coloring scheme with Geneious V 10.2.6

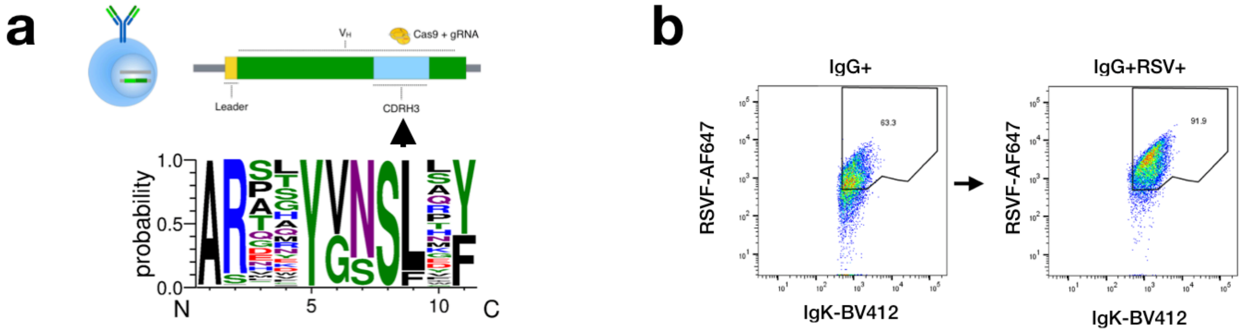


**Supplementary Figure 7. RSV3 CDRH3 antibody library screening workflow.**
a, RSV3 CDRH3 libraries were generated by CRISPR-Cas9 homology directed mutagenesis using an ssODN with degenerate codons representing a sequence space depicted by the logo shown. b, Transfected cells were subsequently sorted in two consecutive steps for antibody expression and specificity or negativity towards RSV-F.

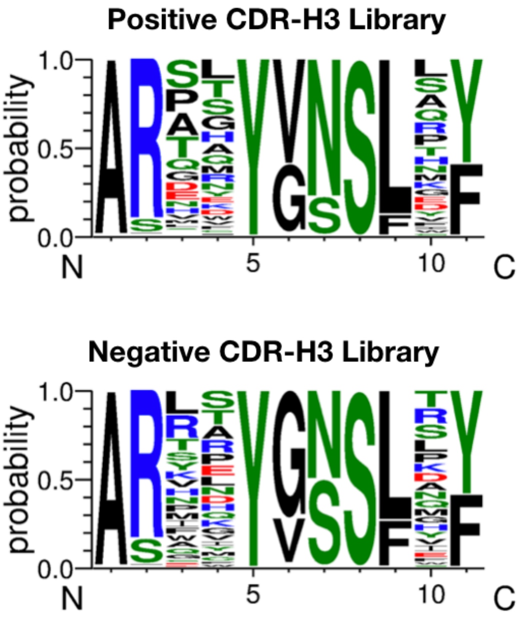


**Supplementary Figure 8. Deep sequencing results of RSV3 CDRH3 library screening.**
Sequence logos show the aggregated sequences found in the a, positive and b, negative fractions of the RSV3 CDRH3 library screen.

| Group | n= | Primary | | 1st Boost | | 2nd Boost | | 3rd Boost | Log $EC_{50}$ |
|---|---|---|---|---|---|---|---|---|---|
| | | *Antigen* | *Adjuvant* | *Antigen* | *Adjuvant* | *Antigen* | *Adjuvant* | *Antigen* | |
| Ova, 3 boosts | 3 | 200 µg Ova | 20 µg MPLA | 50 µg Ova | 20ug MPLA | 50 µg Ova | 20 µg MPLA | 50 µg Ova | 5.19 |
| Ova, 2 boosts | 3 | 200 µg Ova | 20 µg MPLA | 50 µg Ova | 20ug MPLA | 50 µg Ova | | | 5.10 |
| Ova, 1 boost | 3 | 200 µg Ova | 20 µg MPLA | 50 µg Ova | | | | | 4.97 |
| Ova, no boost | 3 | 200 µg Ova | 20 µg MPLA | | | | | | 2.20 |
| HEL, 3 boosts | 3 | 200 µg HEL | 20 µg MPLA | 50 µg HEL | 20ug MPLA | 50 µg HEL | 20 µg MPLA | 50 µg HEL | 4.38 |
| HEL, 2 boosts | 3 | 200 µg HEL | 20 µg MPLA | 50 µg HEL | 20ug MPLA | 50 µg HEL | | | 4.24 |
| HEL, 1 boost | 3 | 200 µg HEL | 20 µg MPLA | 50 µg HEL | | | | | 3.72 |
| HEL, no boost | 3 | 200 µg HEL | 20 µg MPLA | | | | | | 2.33 |
| BCP, 3 boosts | 3 | 200 µg BCP | 20 µg MPLA | 50 µg BCP | 20ug MPLA | 50 µg BCP | 20 µg MPLA | 50 µg BCP | 4.67 |
| BCP, 2 boosts | 3 | 200 µg BCP | 20 µg MPLA | 50 µg BCP | 20ug MPLA | 50 µg BCP | | | 4.42 |
| BCP, 1 boost | 3 | 200 µg BCP | 20 µg MPLA | 50 µg BCP | | | | | 4.19 |
| BCP, no boost | 3 | 200 µg BCP | 20 µg MPLA | | | | | | 2.23 |
| RSV-F, 2 boosts | 10 | 10 µg RSV-F | 1% Alum | 10 µg RSV-F | 1% Alum | 10 µg RSV-F | 1% Alum | | 4.35 |

**Supplementary Table 1. Immunization scheme and antigen cohorts.**
Mice were boosted zero to three times depending on antigen and cohort. MPLA was used as adjuvant for OVA, HEL and BCP injections while Alum was used for RSV-F immunizations. Antigen-specific serum antibody titre as determined by ELISA shown as the $EC_{50}$, which is the fold dilution at which half of the maximum response occurs.

| Clone-ID | HC sequence | LC sequence |
|---|---|---|
| OVA1 | QVQLQQSGAELVRPGSSVKISCKASGYAFSSYWMNWVKQRPGQGLEWIGQINPGDGDTYNNGKFKGKATLTADKSSSTAYMQLNNLTSKDSAVYFCARGGYGSFAYWGQGTLVTVSA | DIVLTQSPAIMSASPGEKVTISCSASSSVSYMFWYQQKPGSSPKPWIYRTSNLASGVPARFSGSGSGTSYSLTISSMEAEDAATYYCQQYHSYPLTFGAGTKLEIK |
| OVA2 | QVQLKQSGAELVRPGASVKLSCKTSGYIFTSYWIHWVKQRSGQGLEWIARIYPGTGSSYYNEKFKGKATLTADKSSSTAYMQLSSLKSEDSAVYFCARAEYDSSYAMDFWGQGTSVTVSS | DIVLTQSPASLAVSLGQRATISCRASQSVSTSSYSFMNWYQQKPGQPPKLLIKSASNLESGVPARFSGSGSGTDFTLNIHPVEEEDTATYYCQHSWEIPLTFGAGTKLEIK |
| OVA3 | QVQLKQSGAELVRPGASVKLSCKTSGYIFTSYWIHWVKQRSGQGLEWIARIYPGTGSTYYNEKFKGKATLTADKSSSTAYMQLSSLKSEDSAVYFCARGDYGSSYFDYWGQGTTLTVSS | DIVMTQSPASLAVSLGQRATISCRASQSVSTSSYSFMHWYQQKPGQPPKLLIKSASNLESGVPARFSGSGSGTDFTLNIHPVEEEDTATYYCQHSWEIPLTFGAGTKLEIK |
| OVA4 | QVQLKQSGAELVRPGASVKLSCKTSGYIFTSYWIHWVKQRSGQGLEWIARIYPGTGSTYYNEKFKGKATLTADKSSSTAYMQLSSLKSEDSAVYFCARNGDYAMDYWGQGTSVTVSS | DIVMTQSPASLAVSLGQRATISCRASQSVSTSSYSFMHWYQQKPGQPPKLLIKYASNLESGVPARFSGSGSGTDFTLNIHPVEEEDTATYYCQHSWEIPYTFGGGTKLELK |
| OVA5 | QVQLQQPGAELVKPGASVKLSCKASGYTFTSYYMYWVKQRPGQGLEWIGEINPSNGGTNFNEKFKSKATLTVDKSSSTAYMQLSSLTSEDSAVYYCTRGGYGSFAYWGQGTLVTVSA | DILMTQSPAIMSASPGEKVTISCSASSSVSYMFWYQQKPGSSPKPWIYRTSNLASGVPARFSGSGSGTSYSLTISSMEAEDAATYYCQQYHSYPLTFGSGTKLEIK |
| OVA6 | QVQLQQSGAELVRPGASVKLSCKTSGYIFTSYWIHWVKQRSGQGLEWIARIYPGTGSTYYNEKFKGKATLTADKSSSTAYMQLSSLKSEDSAVYFCAITTVNAMDYWGQGTSVTVSS | DIQMTQSPASLAVSLGQRATISCRASQSVSTSSYSHMNWYQQKPGQPPKLLIKYASNLESGVPARFSGSGSGTDFTLNIHPVEEEDTAIYYCQHSWEMPYTFGGGTKLEIK |

**Supplementary Table 2. Sequences confirmed to bind OVA.**

| | Surrogate LC sequences |
|---|---|
| OVA1 | DIVMTQSPAIMSASPGEKVTMTCSASSSVSYMHWYQQKSSTSPKLWIYDTSKLPSGVPGRFSGSGSGNSYSLTISSMEAEDVATYYCFQGSGYPLTFGAGTKLEIK |
| OVA5 | DIVMTQSPSSMYASLGERVTITCKASQDIKSYLSWFQQKPGKSPKTLIYRANRLVDGVPSRFSGSGSGQDYSLTISSLEYEDMGIYYCLQYDEFPYTFGGGTKLEIK |
| | DIQMTQSPSSMYASLGERVTITCKASQDINSYLSWFQQKPGKSPKTLIYRANRLVDGVPSRFSGSGSGQDYSLTISSLEYEDMGIYYCLQYDEFPYTFGGGTKLEIK |
| | DIVLTQSHKFMSTSVGDRVSITCKASQDVGSAVAWYQQKPGQSPKLLIYWASTRRTGVPDRFTGSGSGTDFTLTINNVQSEDLADYFCHQYNTYSLTFGAGTKLEIK |
| | DIVMTQSPAIMSASPGEKVTMTCSASSSVSYMYWYQQKPGSSPRLLIYDTSNLASGVPVRFSGSGSGTSYSLTISRMEAEDAATYYCQQWSSYPPTFGSGTKLEIK |
| | DIVITQSPAIMSASPGEKVTMTCSASSSVSYMHWYQQKSSTSPKLWIYDTSKLVSGVPGRFSGSGSGNSYSLTISSMEAEDVATYYCFQGSGYPLTFGAGTKLEIK |
| | DIVLTQSPAIMSASPGEKVTMTCSASSSVRYMCWYQQKSSTSPKLWIYDTSKLASGVPGRFSGSGSGNSYSLTIGSMEAEDVATYYCFQGSGYPLTFGAGTKLELK |

**Supplementary Table 3. Surrogate $V_L$ chain sequences for OVA1, OVA5.**

| | HC sequence | LC sequence |
|---|---|---|
| RSV1 | EVQLQQSGAELVRSGASVKLSCTASGFNIKDYYIHWVKQRPEQGLEWIGWIDPENGDTEYAPKFQDKATMTADTSSNTAYLQLSSLTSGDTAVYYCNAWDGNLDYWGQGTSVTVSS | DIQMTQTPLSLPVSLGNQASISCRSSQSIVHSDGNTYLEWYLQKPGQSPKLLIYKVSNRFSGVPDRFSGSGSGTDFTLKISRVEAEDLGVYYCFQGSHVPRTFGGGTKLEIK |
| RSV2 | QVTLKESGPGILQPSQTLSLTCSFSGFSLSTSGMGVSWIRQPSGKGLEWLAHIYWDDDKRYNPSLKSRLTISKDTSRNQVFLKITSVDTADTATYYCARRSYGSSLDYWGQGTTLTVSS | DIVMTQTTSSLSASLGDRVTISCRSSQDISNYLNWYQQKPDGTVKLLIYYTSRLHSGVPSRFSGSGSGSDYSLTITNLEQEDIATYFCQQGDTLPYTFGGGTKLEIK |
| RSV3 | QVQLKESGPGLVAPSQSLSITCTVSGFSLTDYGVNWVRQPPGKGLEWLGMIWGDGSTDYNSALKSRLSISKDISKSQVFLKMNSLQTDDTAKYYCARGNYGNSLDYWGQGTTLTVSS | DIVMTQSPASQSASLGESVTITCLASQTIGKWLAWYQQKPGKSPQLLIYAATILADGVPSRFSGSGSGTKFSFKISSLQAEDFVSYYCQQLYSTPLTFGGGTKLEIK |

**Supplementary Table 4. Sequences confirmed to bind RSV.**

| | Surrogate LC sequences |
|---|---|
| RSV1 | DILMTQSPLSLPVSLGDQVSISCRSSQSIVHSNGNTYLEWYLQKPGQSPKLLIYKVSNRFSGVPDRFSGSGSGTDFTLKISRVEAEDLGVYYCFQGSHVPRTFGGGTKLEIK |
| RSV2 | DIVMTQTTSSLSASLGDRVTISCRASQDISNYLNWYQQKPDGTVKLLIYYTSRLHSGVPSRFSGSGSGTDYSLTISNLEQEDIATYFCQQGNTLPWTFGGGTKLELK |
| RSV3 | DIQINQSPASQSASLGESVTITCLASQTIGKWLAWYQQKPGKSPQLLIYAATSLADGVPSRFSGSGSGTKFSFKISSLQAEDFVSYYCQQLYSTPLTFGGGTKLEIK |

**Supplementary Table 5. Surrogate $V_L$ chain sequences for RSV1, 2 and 3.**

| Target/Antigen | Working concentration | Dilution from stock | Incubation volume | Fluorophore | Product ID |
|---|---|---|---|---|---|
| anti-IgG2c | 13 g/ml | 1:100 | 100 µl | AlexaFluor 488 | 115-545-208 (Jackson ImmunoResearch) |
| anti-IgK | 2.5 g/ml | 1:80 | 100 µl | Brilliant Violet 421 | 409511 (BioLegend) |
| Hen egg lysozyme | 0.99 g/ml | 1:50 | 100 µl | AlexaFluor 647 | 62971-10G-F (Sigma) |
| Ovalbumine | 1.5 g/ml | 1:50 | 100 µl | AlexaFluor 647 | A5503 (Sigma) |
| Blue carrier protein | 1.2 g/ml | 1:50 | 100 µl | AlexaFluor 647 | 77130 (Thermo) |
| RSVF-DS2-biotin | 14 g/ml | 1:50 | 100 µl | - | - |
| Streptavidin | 5 g/ml | 1:100 | 100 µl | AlexaFluor 647 | 405237 (BioLegend) |

**Supplementary Table 6. Flow cytometry labeling components with their respective working concentrations.** HEL, OVA and BCP were labeled in-house with the AlexaFluor 647 Protein Labeling Kit (Thermo, A20173).

| Oligonucleotide ID | Oligonucleotide sequence (5' -> 3') |
|---|---|
| crRNA-JP | GUCAUGGAAGGUUCGGUCAAGUUUUAGAGCUAUGCU |
| crRNA DN_RSV7_H3-3 | GCCUUGGCCCCAGUAGUCAAGUUUUAGAGCUAUGCU |
| DN_RSV3-OOF ssODN | GAACAGTCTGCAAACTGATGACACAGCCAAATATTATTGTGCACGTTAGA |
| ssODN library sequence | CCATTCTTACCTGAGGAGACTGTGAGAGTGGTTCCCTGGCCCCARWAM |
| FW primer PCR1 | CCCTCCTTTAATTCCCCAGCTCTCAAATCCAGACTG |
| REV primer PCR1 | GAGGAGAGAGAGAGAGTGCAATGCTCAGAAAACTCC |
| cr_DM1 | TTGTGCACGTTAGAGCCAGGCGG |
| DN_VHscreenSF_GA_PnPs1_fw | CCTTCCGGGGATCCTG |
| DN_VHscreenSF_GA_PnPs1_rev | CCTGGAGAGGCCATTCTTAC |
| CP399 | AAGAATGGCCTCTCCAGGTCTTTATTTTTAAC |
| CP400 | TGACAGGATCCCCGGAAGG |
| SF_062 | CACACSASTGWGGC |
| DN_LC_lib_GA_fw | GGCCCCAACTGTATCCAT |
| DN_LC_lib_GA_rev | CCGGGACATTATAACTGAAGC |
| DN_MLC_V_GA_fw1 | GCTTCAGTTATAATGTCCCGGGGGGAYATCCAGCTGACTCAGCC |
| DN_MLC_V_GA_fw2 | GCTTCAGTTATAATGTCCCGGGGGGAYATTGTTCTCWCCCAGTC |
| DN_MLC_V_GA_fw3 | GCTTCAGTTATAATGTCCCGGGGGGAYATTGTGMTMACTCAGTC |
| DN_MLC_V_GA_fw4 | GCTTCAGTTATAATGTCCCGGGGGGAYATTGTGYTRACACAGTC |
| DN_MLC_V_GA_fw5 | GCTTCAGTTATAATGTCCCGGGGGGAYATTGTRATGACMCAGTC |
| DN_MLC_V_GA_fw6 | GCTTCAGTTATAATGTCCCGGGGGGAYATTMAGATRAMCCAGTC |
| DN_MLC_V_GA_fw7 | GCTTCAGTTATAATGTCCCGGGGGGAYATTCAGATGAYDCAGTC |
| DN_MLC_V_GA_fw8 | GCTTCAGTTATAATGTCCCGGGGGGAYATYCAGATGACACAGAC |
| DN_MLC_V_GA_fw9 | GCTTCAGTTATAATGTCCCGGGGGGAYATTGTTCTCAWCCAGTC |
| DN_MLC_V_GA_fw10 | GCTTCAGTTATAATGTCCCGGGGGGAYATTGWGCTSACCCAATC |
| DN_MLC_V_GA_fw11 | GCTTCAGTTATAATGTCCCGGGGGGAYATTSTRATGACCCARTC |
| DN_MLC_V_GA_fw12 | GCTTCAGTTATAATGTCCCGGGGGGAYRTTKTGATGACCCARAC |
| DN_MLC_V_GA_fw13 | GCTTCAGTTATAATGTCCCGGGGGGAYATTGTGATGCBCAGKC |
| DN_MLC_V_GA_fw14 | GCTTCAGTTATAATGTCCCGGGGGGAYATTGTGATAACYCAGGA |
| DN_MLC_V_GA_fw15 | GCTTCAGTTATAATGTCCCGGGGGGAYATTGTGATGACCCAGWT |
| DN_MLC_V_GA_fw16 | GCTTCAGTTATAATGTCCCGGGGGGAYATTGTGATGACACAACC |
| DN_MLC_V_GA_fw17 | GCTTCAGTTATAATGTCCCGGGGGGAYATTTTGCTGACTCAGTC |
| DN_MLC_V_GA_fw18 | GCTTCAGTTATAATGTCCCGGGGGGARGCTGTTGTGACTCAGGAATC |
| DN_MLC_J_GA_rev1 | ATGGATACAGTTGGGGCCGCGTCGGCCCGTTTGATTTCCAGCTTGG |
| DN_MLC_J_GA_rev2 | ATGGATACAGTTGGGGCCGCGTCGGCCCGTTTTATTTCCAGCTTGG |
| DN_MLC_J_GA_rev3 | ATGGATACAGTTGGGGCCGCGTCGGCCCGTTTTATTTCCAACTTTG |
| DN_MLC_J_GA_rev4 | ATGGATACAGTTGGGGCCGCGTCGGCCCGTTTCAGCTCCAGCTTGG |
| DN_MLC_J_GA_rev5 | ATGGATACAGTTGGGGCCGCGTCGGCCCGTAGGACAGTCAGTTTGG |
| DN_MLC_J_GA_rev6 | ATGGATACAGTTGGGGCCGCGTCGGCCCGTAGGACAGTGACCTTGG |
| CP467 | CATGTGCCTTTTCAGTGCTTTCTC |
| CP468 | CTAGATGCCTTTCTCCCTTGACTC |
| CP99 | GAAAACAACATATGACTCCTGTCTTC |
| CP168 | TGACCTTCTCAAGTTGGC |
| DM125 | AATGATACGGCGACCACCGAGATCTACACTCTTTCCCTACACGACGCTCT |
| DM142 | CAAGCAGAAGACGGCATACGAGATGTGCGGACGTGACTGGAGTTCAGA |
| DM143 | CAAGCAGAAGACGGCATACGAGATCGTTTCACGTGACTGGAGTTCAGAC |
| DM144 | CAAGCAGAAGACGGCATACGAGATAAGGCCACGTGACTGGAGTTCAGA |

**Supplementary Table 7. Oligonucleotides used in this study.**
